# Human salivary amylase gene copy number impacts oral and gut microbiomes

**DOI:** 10.1101/359737

**Authors:** Angela C. Poole, Julia K. Goodrich, Nicholas D. Youngblut, Guillermo G. Luque, Albane Ruaud, Jessica L. Sutter, Jillian L. Waters, Qiaojuan Shi, Mohamed El-Hadidi, Lynn M. Johnson, Haim Y. Bar, Daniel H. Huson, James G. Booth, Ruth E. Ley

## Abstract

Host genetic variation influences the composition of the human microbiome. While studies have focused on associations between the microbiome and single nucleotide polymorphisms in genes, their copy number (CN) can also vary. Here, in a study of human subjects including a 2-week standard diet, we relate oral and gut microbiome to CN at the *AMY1* locus, which encodes the gene for salivary amylase, active in starch degradation. We show that although diet standardization drove gut microbiome convergence, *AMY1*-CN influenced oral and gut microbiome composition and function. The gut microbiomes of low-*AMY1*-CN subjects had an enhanced capacity for breakdown of complex carbohydrates. Those of high-*AMY1* subjects were enriched in microbiota linked to resistant starch fermentation, had higher fecal SCFAs, and drove higher adiposity when transferred to germfree mice. Gut microbiota results were validated in a larger separate population. This study establishes *AMY1*-CN as a genetic factor patterning microbiome composition and function.

## Introduction

Host genotype has recently emerged as a significant factor determining the relative abundance of specific members of the human microbiota (Goodrich et al. 2016; Rothschild et al. 2018). Heritable gut microbes, whose abundances are influenced by host genotype, have been identified, and genome-wide association studies (GWAS) have linked specific gene variants to members or functions of the gut microbiome (Goodrich et al. 2017; Lim et al. 2017; Davenport 2016; Turpin et al. 2016; Davenport et al. 2015; Goodrich et al. 2014; Goodrich et al. 2016). In addition to allelic differences between individuals, another important aspect of human genetic variation is the copy number (CN) of genes. Gene duplications resulting in increased CN provide a rapid means of adaptation to environmental change (Iskow et al. 2012). Copy number variation (CNV) in genes accounts for far more genomic variability than SNPs (Conrad et al. 2010) and can influence a significant amount of gene expression (Chiang et al. 2017). This important aspect of human genetic variability likely relates to microbiome differences between individuals, but links between the CNV of specific human genes and the microbiome remain to be elucidated.

CN variation of the *AMY1* gene, which encodes the human salivary amylase enzyme, is considered one of the strongest signals of recent selection on human populations. Salivary amylase hydrolyses alpha bonds of starch and glycogen, beginning the process of starch degradation in the mouth. A dietary shift to greater starch intake during the agronomic transition of the Neolithic period is thought to have selected for the duplications observed within the salivary amylase gene locus (Kelley and Swanson 2008; Iskow et al. 2012; Perry et al. 2015). Today, the mean *AMY1*-CN is reported higher in populations with an agrarian background compared to hunter-gatherers (Perry et al. 2007). Across genetic backgrounds, human *AMY1*-CN ranges from 2 to 24 (Perry et al. 2007; Usher et al. 2015; Yong et al. 2016).

Because *AMY1*-CN is positively correlated with amylase activity in the mouth (Mandel et al. 2010; Perry et al. 2007), it has the potential to influence carbohydrate processing and gut microbiome composition. Complex carbohydrates include polysaccharides, such as starch, that are digested by host amylases and by gut microbial carbohydrate-active enzymes. These carbohydrates encounter host amylase first in the mouth, then again after passing the stomach, where pancreatic amylase is added to the chyme, and the liberated sugars are absorbed in the small intestine (SI). Uptake of sugars liberated by host enzymes from starch in the SI yields more energy to the host than does uptake in the large intestine (LI) of microbial products of starch fermentation (Walter and Ley 2011). The resulting host-microbial competition for energy-rich starch may have driven selection for duplications at the amylase locus. Indeed, amylase supplementation to farm animals (AMY1L) enhances starch digestibility and promotes weight gain (Burnett 1962; Gracia et al. 2003; Jo et al. 2012). Similarly, humans with a high *AMY1*-CN (AMY1H), who produce high levels of salivary amylase, should derive more energy from the same carbohydrate-rich diet than those with a low *AMY1*-CN (AMY1L). Compared to AMY1H, AMY1L individuals might be expected to harbor gut microbiomes with a greater capacity for breakdown of complex carbohydrates, compensating for the lower levels of host amylase.

Due to its link to carbohydrate digestion, *AMY1*-CN has been investigated for associations with BMI and metabolism. Results of these studies have been somewhat inconsistent across populations, with low *AMY1*-CN associated with high BMI in some populations (Viljakainen et al. 2015; Mejía-Benítez et al. 2015; Falchi et al. 2014; Marcovecchio et al. 2016; Bonnefond et al. 2017), but not others (Usher et al. 2015; Yong et al. 2016). Part of the discrepancy between outcomes may be methodological (Usher et al. 2015). Starch intake may also be an important variable in the relationship between *AMY1*-CN and BMI (Rukh et al. 2017). Furthermore, the gut microbiome, which is known to interact with host genetics (Blekhman et al. 2015; Goodrich et al. 2014; Bonder et al. 2016; Wang et al. 2016), and to impact host metabolism (Sonnenburg and Bäckhed 2016; Zeevi et al. 2015; Pedersen et al. 2016; Goodrich et al. 2014), may also interact with host *AMY1*-CN to affect phenotypes. Whether host *AMY1*-CN status regulates the composition and function of the gut microbiomes remains to be ascertained.

Here, we address the question of how *AMY1*-CN relates to diversity and function of the gut microbiomes of healthy individuals with normal BMIs. From a group of >100 volunteers for whom we quantified *AMY1*-CN, we recruited 25 participants into a one-month longitudinal study. To standardize diets and ensure starch consumption by all participants, we provided all meals during the middle 2 weeks of the study, and participants kept a food diary for the duration. Subjects provided oral and stool samples three times weekly, and we used 16S rRNA gene sequence analysis to assess the effects of host *AMY1*-CN and diet intervention on oral and fecal microbiomes. We then used a larger separate population to validate the results from the gut microbiome dataset. In addition, we obtained a functional characterization of fecal microbiomes through (i) deep metagenomic sequencing, (ii) short chain fatty acid (SCFA) measures, and (iii) fecal transfers to germfree mice.

## Results

### Range of *AMY1*-CN in study participants

We collected buccal swabs from 105 volunteers recruited on the Cornell University campus and measured their *AMY1*-CN by qPCR (Figure 1A, Table S1), then selected 25 participants across the *AMY1*-CN distribution for further study. We confirmed the *AMY1*-CN of participants using alternate qPCR primers and digital PCR (See STAR Methods, Table 1). Based on the results, 11 participants were assigned to a high group (CN > 8, designated AMY1H), 5 to a medium (5 < CN < 8, AMY1M) and 9 to a low group (CN < 5, AMY1L). Neither BMI nor body fat percentage differed significantly between groups (Table 1). Pancreatic amylase, *AMY2*, had a smaller CN range than *AMY1*, and these measures were positively correlated (Spearman’s rho = 0.79, p = 3×10^−8^; Table 1).

**Figure 1.**
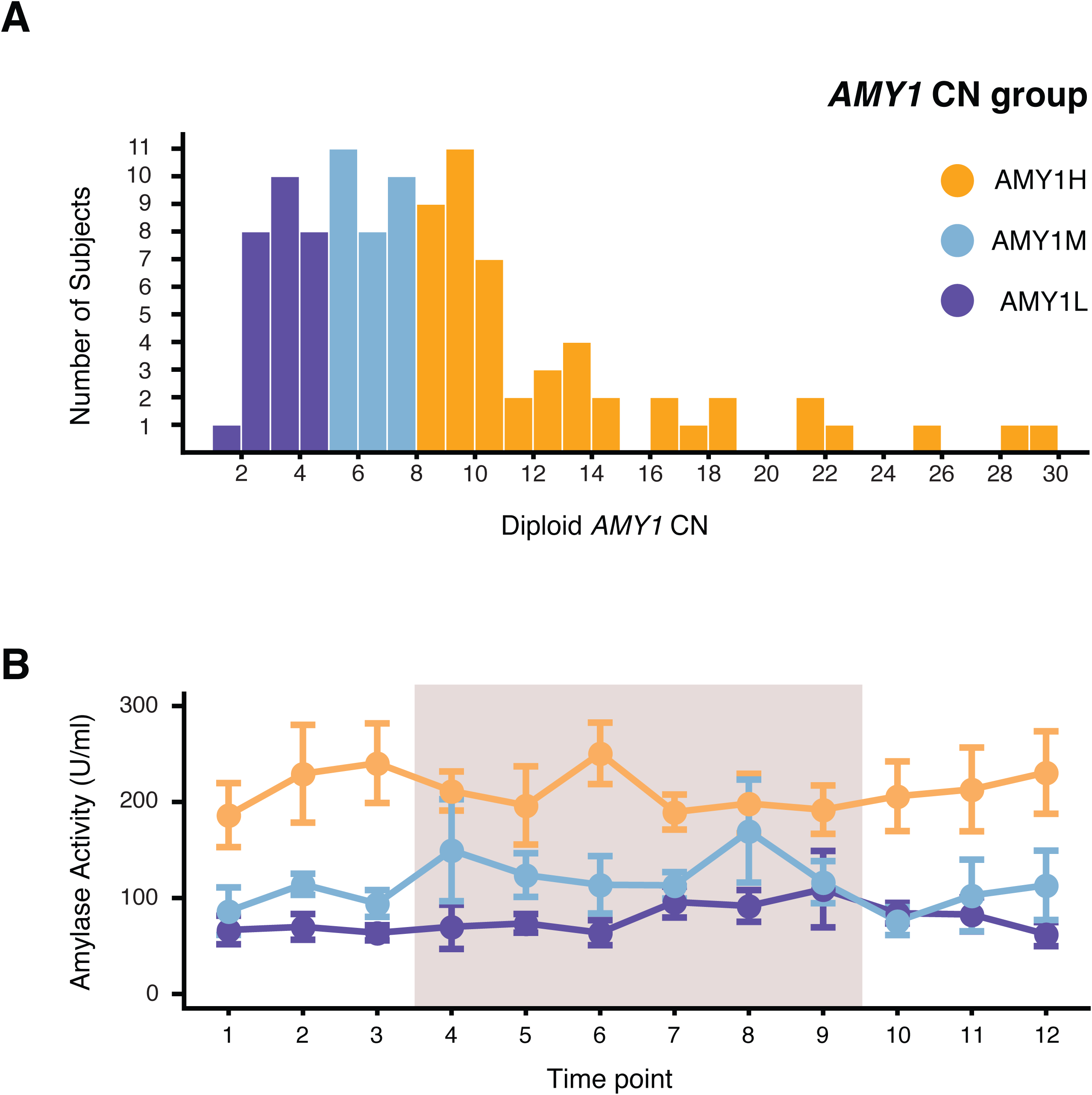
*AMY1*-CN distribution for 105 subjects recruited at Cornell University. (A) Diploid *AMY1*-CN distribution for 105 subjects was obtained using qPCR with primer sequences previously reported by (Perry et al. 2007). (B) Mean amylase activity per ml of saliva ± SEM for the 3 AMY1 groups. Measurements were performed in triplicate for both qPCR and salivary amylase activity.

**Table 1.**
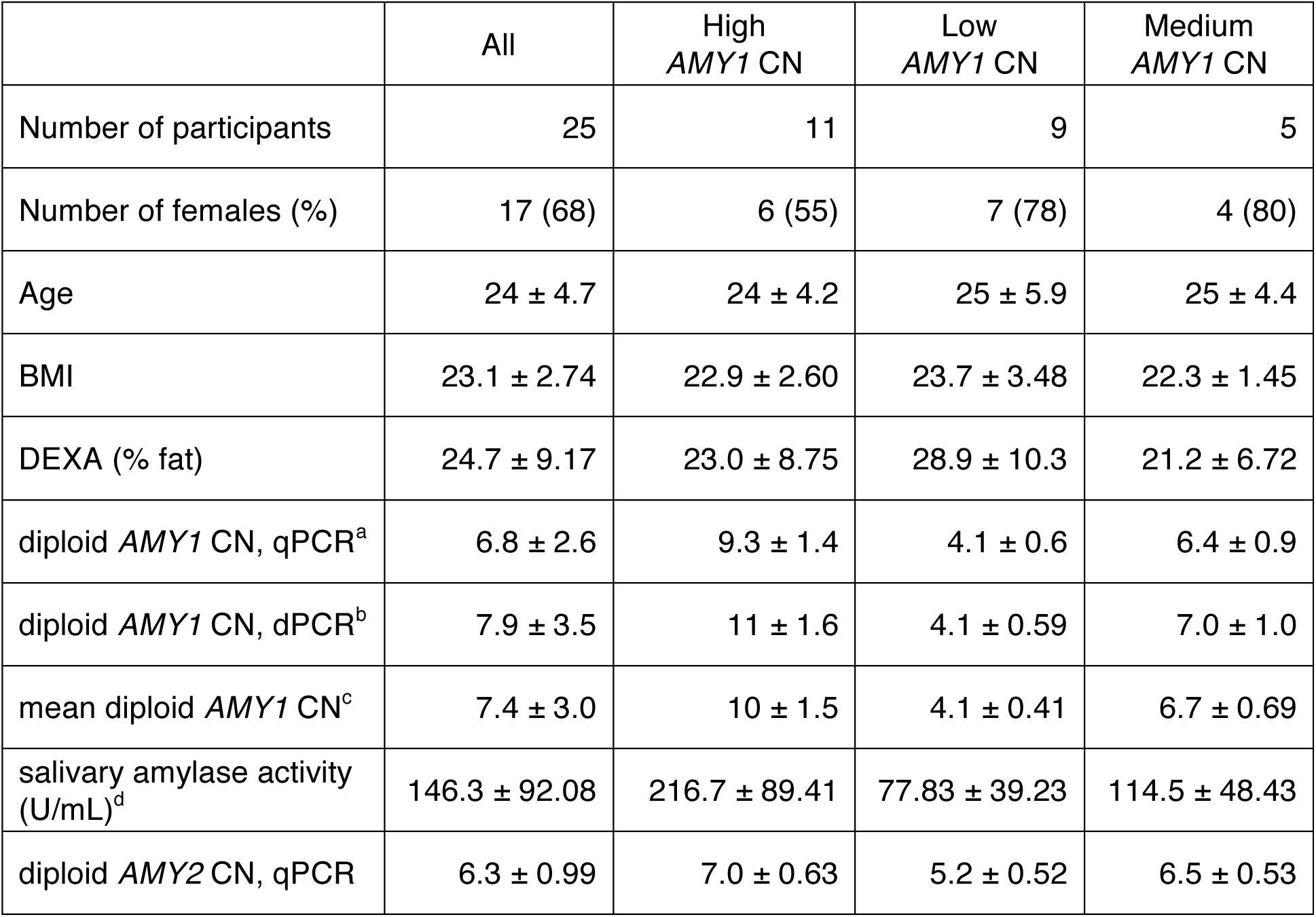
Characteristics of study participants. Diploid *AMY1* CN was confirmed using qPCR^a^and digital PCR^b^. The mean diploid *AMY1* CN^c^is the mean of both measurements and was used in subsequent statistical correlations. *AMY1* CN group was determined by the following criteria: AMY1H = mean diploid *AMY1* CN > 8, AMY1L = mean diploid *AMY1* CN < 5. For all measurements with ranges, mean ± standard deviation is provided. Salivary amylase activity was measured for all samples collected (12 TPs for all participants) and for each participant, the mean salivary amylase activity across all TPs was calculated. For statistical analyses using ordinal categories, the medium *AMY1* CN group was excluded. See also Tables S1 and S2.

### Dietary intake was similar between *AMY1* CN groups throughout the study

To mitigate the effects of diet differences between individuals on their microbiomes and to promote frequent consumption of starch, during the middle two weeks of the study we provided participants with all meals and snacks. Participants ate from the menu freely, occasionally supplemented it (Table S2), and recorded their dietary intake in food diaries (see STAR Methods). Based on these records, mean percentages of total carbohydrate, protein, and fat intake did not differ significantly between the AMY1H, AMY1M, and AMY1L groups throughout the study, regardless of whether meals were consumed during (weeks 2-3) or outside (weeks 1, 4) of the standardized diet period (Figure S1A-C). The intake of all three macronutrients differed between days (p < 1×0^−5^), however, the standardized diet period did not significantly impact mean intake of macronutrients.

### *AMY1* CN versus oral and fecal amylase activity

Saliva samples were obtained at 12 time points (TPs). Amylase enzyme activity in saliva samples ranged between 10.2 - 527 Units per ml of saliva, similar to previously reported ranges (Mandel et al. 2010). *AMY1*-CN correlated with salivary amylase activity (SAA) levels across all subjects at all TPs (linear mixed model; p = 2.1×0^−5^). SAA levels for the AMY1H were higher than for the AMY1L at all TPs (linear mixed model; p = 1.9×0^−4^; Figure 1B), with AMY1M individuals intermediate. Fecal amylase activity (FAA) measured in stool at all TPs was variable within and between subjects (0.6 to 1,120 Ug^−1^ stool); consistent with published reports (Macfarlane and Englyst 1986; Moriyoshi et al. 1991)(Figure S1D), and unlike SAA, did not correlate with *AMY1*-CN. To further characterize FAA, we used an ELISA method to measure enzyme levels of host pancreatic amylase in stool samples at TPs 6 and 10. Across all 25 subjects, pancreatic amylase levels were highly correlated with FAA (Spearman’s rho = 0.80, p = 3.7×0^−6^ for TP 6 and Spearman’s rho = 0.78, p = 6.3×0^−5^ for TP 10; Figure S1E), although *AMY2*-CN was not. This finding corroborates previous reports that FAA is largely pancreatic (Macfarlane and Englyst 1986; Moriyoshi et al. 1991).

### Patterning of the oral microbiota by host *AMY1* CN

Saliva samples were profiled for microbial community diversity by Illumina sequencing of the V4 region of 16S rRNA gene PCR amplicons (Table S1). Sequences were clustered into operational taxonomic units (OTUs) using a threshold of 97% pairwise sequence identity (see STAR Methods). We observed that oral microbiome richness (alpha-diversity) was correlated with *AMY1*-CN (using Chao 1, Observed Species, and Faith’s PD metrics p < 0.01; but not Shannon’s Index), and alpha-diversity was higher in AMY1H than AMY1L individuals (p = 0.011; Figure 2A). Principal coordinates analysis (PCoA) of unweighted UniFrac distance metrics revealed clustering of saliva microbiomes by subject, and a trend for separation by *AMY1*-CN group across all subjects (Figure 2B, S2A, S2B). Together, these observations indicate that across the *AMY1*-CN gradient, a higher *AMY1*-CN associated with greater richness of the microbiome without a significant shift in overall diversity.

**Figure 2.**
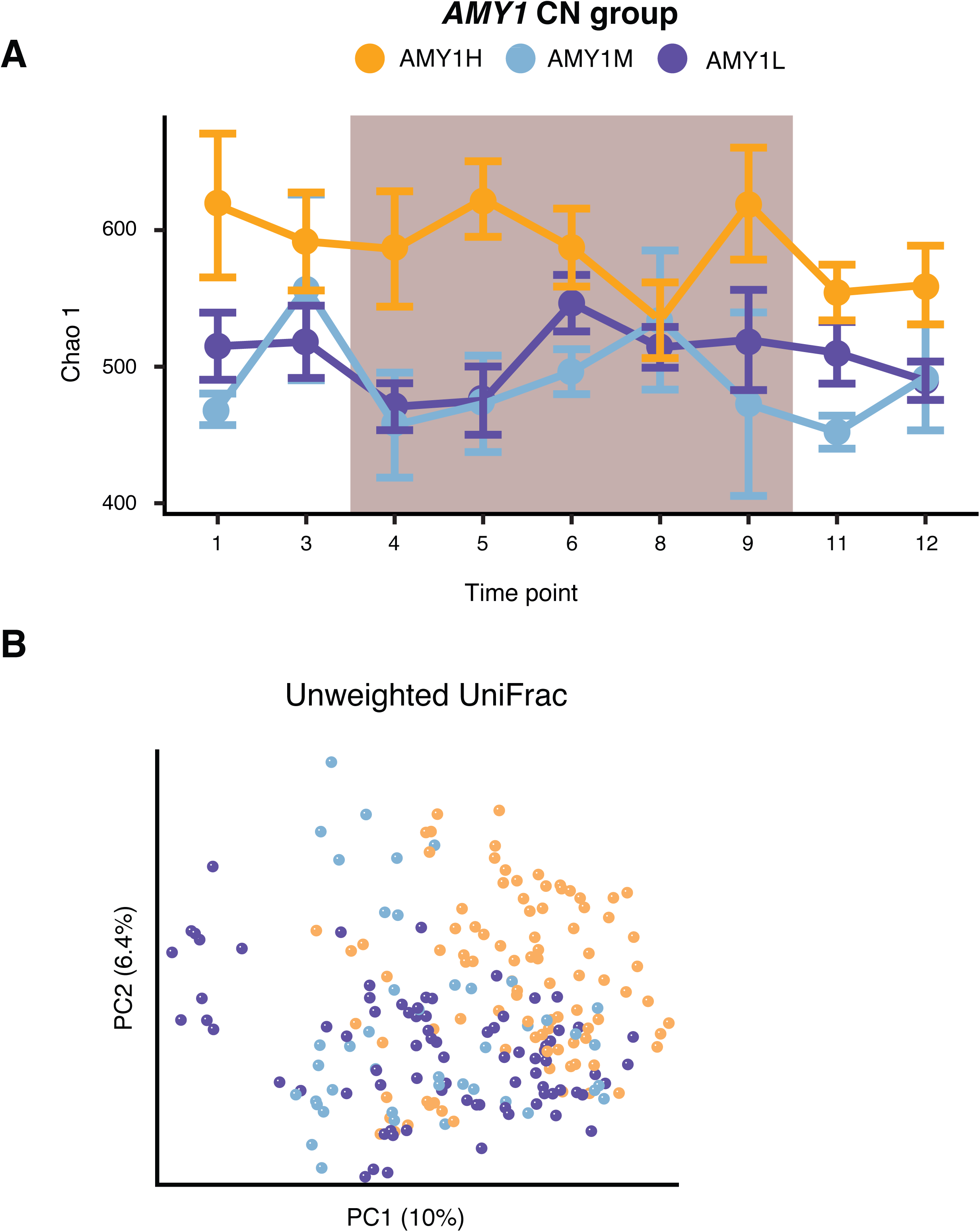
Oral microbiomes differ in diversity between *AMY1*-CN groups. (A) Alpha-diversity assessed with the metric Chao 1. (B) Principal coordinates analysis (PCoA) of the unweighted UniFrac distances between samples collected from all subjects throughout the study. The first two PCs are plotted. The percent variation explained by each PC is indicated on the axes. Samples are colored according to *AMY1*-CN group.

We searched for OTUs that could, based on their relative abundances, distinguish between AMY1H and AMY1L subjects over the whole time period and including all subjects, using a machine learning technique (random forests, STAR Methods). In this approach, a model trained on the 80% of the samples produced an accuracy of 97.67% with a Matthews correlation coefficient (MCC) of 94.92% and an area under the curve (AUC) of 96.66% once it was tested on the remaining 20% of the samples. We then used a feature selection process (STAR methods) to identify the relevant features of the model. Among the top OTUs that most discriminated AMY1H and AMY1L groups (AMY1M excluded) were OTUs classified as *Prevotella* and *Porphyromonas* (Figure 3A). An analysis using all data (AMY1M included, with the *AMY1*-CN classified into high or low by k-means clustering), yielded similar results (Figure S3A).

**Figure 3.**
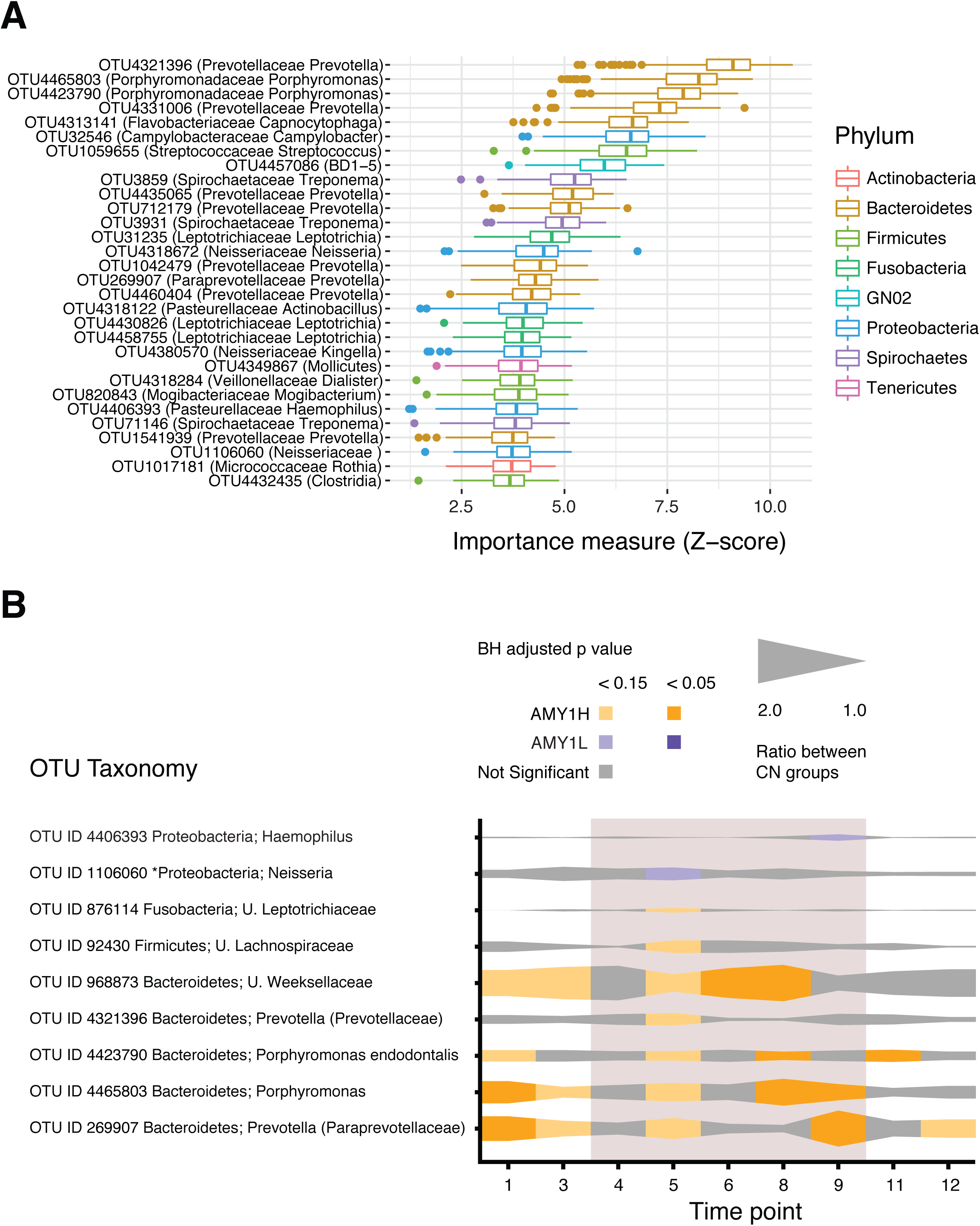
Oral microbiomes differ between SAA and *AMY1*-CN groups at the OTU level. (A) The OTUs shown here were identified using machine learning as distinguishing between SAA groups. The length of the bar represents the magnitude of the mean decrease in Gini index and the orientation indicates the group in which the OTU is enriched. (B) The mean relative abundances of the OTUs included in this plot were significantly different between AMY1H and AMY1L groups at one or more TPs using the statistical model Harvest. Each ribbon corresponds to a single OTU with taxonomy indicated to the left, with unclassified abbreviated “U.” Taxonomy may be shared by several OTUs. The width of the ribbon at each TP shows the ratio of the mean OTU relative abundances between the *AMY1* CN groups. If the ribbon is colored orange at a given TP, the AMY1H group has a higher mean relative abundance of the OTU; when purple, the AMY1L has a higher mean relative abundance. When the ribbon is colored grey, the Benjamini-Hochberg (BH) adjusted p ≥ 0.15. Lighter orange or purple corresponds to a BH adjusted p < 0.15; darker colors correspond to BH adjusted p < 0.05. * The asterisk denotes an OTU that was assigned taxonomy with higher resolution after performing a BLAST search using the representative sequence. See also Table S3.

To gain a time-resolved view into the taxa driving differences for subjects at the extremes of the *AMY1* CN gradient, we compared AMY1H and AMY1L groups (AMY1M excluded) at each time point using a bivariate model (hereafter referred to as ‘Harvest’) (Bar, Booth, and Wells 2014). This analysis revealed a total of 9 OTUs with significantly different mean relative abundances between AMY1H and AMY1L (both Benjamini-Hochberg (BH) adjusted p < 0.05 and BH adjusted p < 0.15 are shown; Figure 3B); none exhibited different variances. As observed for the machine-learning analysis, OTUs that discriminated the AMY1H and AMY1L groups belonged to the genera *Prevotella*, *Haemophilus*, *Neisseria*, and *Porphyromonas* (Fig. 3B). These patterns highlight that (i) OTUs are consistently elevated in either AMY1H or AMY1L, and the direction stays constant over time, and (ii) their enrichment in one group or another moves in and out of statistical significance over time.

### Patterning of the fecal microbiota by host *AMY1* CN

In contrast to oral microbiomes, the richness of fecal microbiomes was generally similar between *AMY1*-CN groups (using Chao 1, Observed Species, and Faith’s PD metrics; Figure 4A). UniFrac analyses showed that overall microbial diversity was unrelated to host *AMY1*-CN (Figure 4B, S4A), with some clustering by subject (Figure S4B). Using a random forest analysis as above, the prediction performed on the 20% of the samples reserved as testing dataset produced an accuracy of 98.21% with an MCC of 96.04% and an AUC of 97.36%. After performing a feature selection process, we identified OTUs discriminating between AMY1H and AMY1L (AMY1M excluded) as belonging to the Ruminococcaceae (*Ruminococcus* and *Oscillospira*), and Lachnospiraceae (*Blautia*, *Dorea*, *Roseburia*; Figure 5A; Table S4). Similarly, when all 25 subjects were reclassified into low and high groups based on k-means clustering (AMY1M included, as above), a similar set of discriminatory OTUs was observed (Figure S5). We also identified taxa that differentiated AMY1H and AMY1L groups at each TP using Harvest): 7 of the 11 discriminatory OTUs were members of the Ruminococcaceae family, and all but one were elevated in the AMY1H compared to the AMY1L group (Figure 5B). As observed in the oral microbiota, the Harvest analysis showed that discriminatory OTUs were consistently enriched in the same *AMY1*-CN category, although the significance of the enrichment could vary with TP. Members of the Ruminococcaceae have been linked to resistant starch degradation; enrichment of Ruminococcaceae OTUs in the AMY1H is consistent with reduced availability of starches susceptible to host amylase degradation in the distal gut of the AMY1H host.

**Figure 4.**
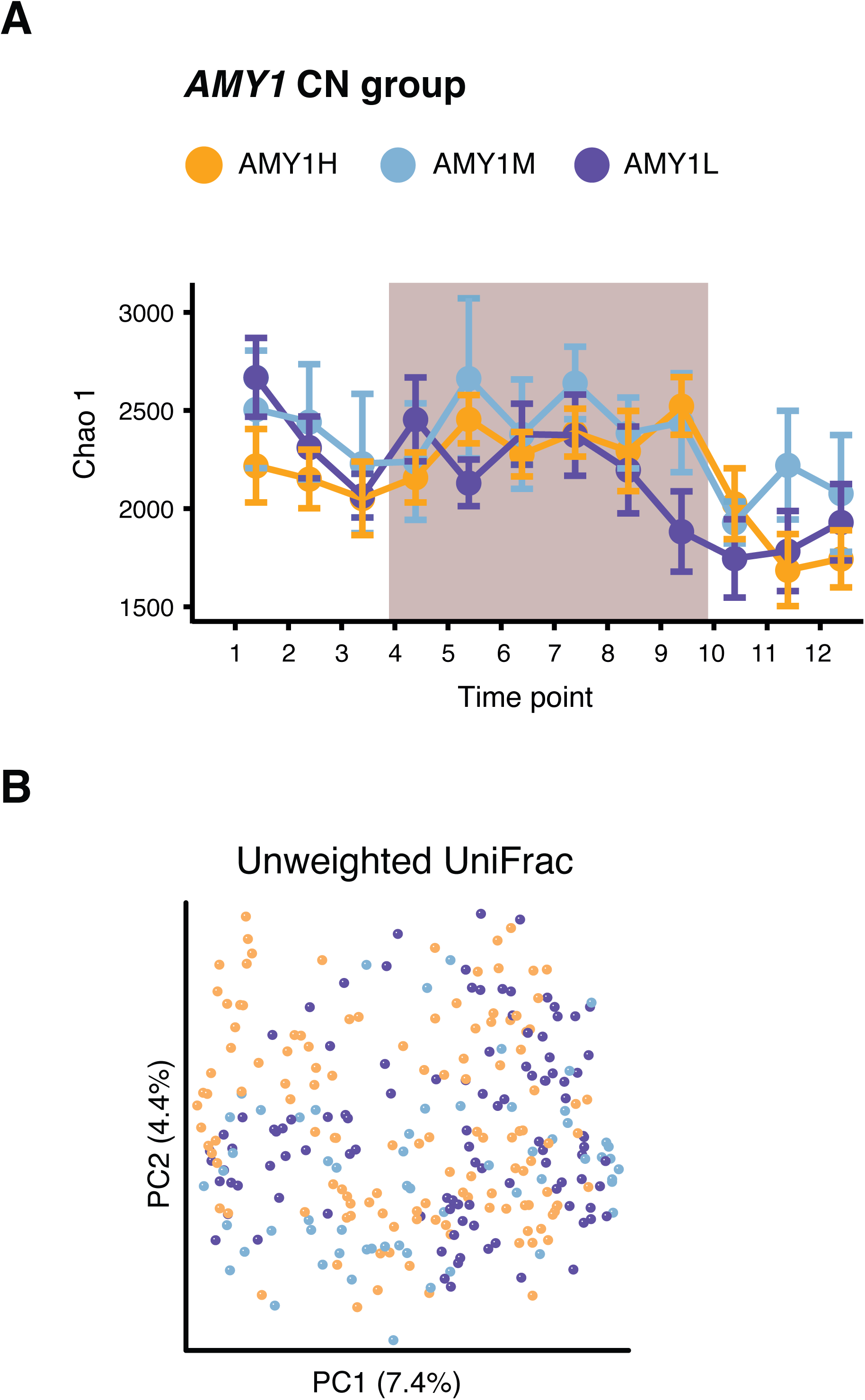
Gut microbiomes do not differ in overall composition between *AMY1* CN groups. See legend for Figure 2.

**Figure 5.**
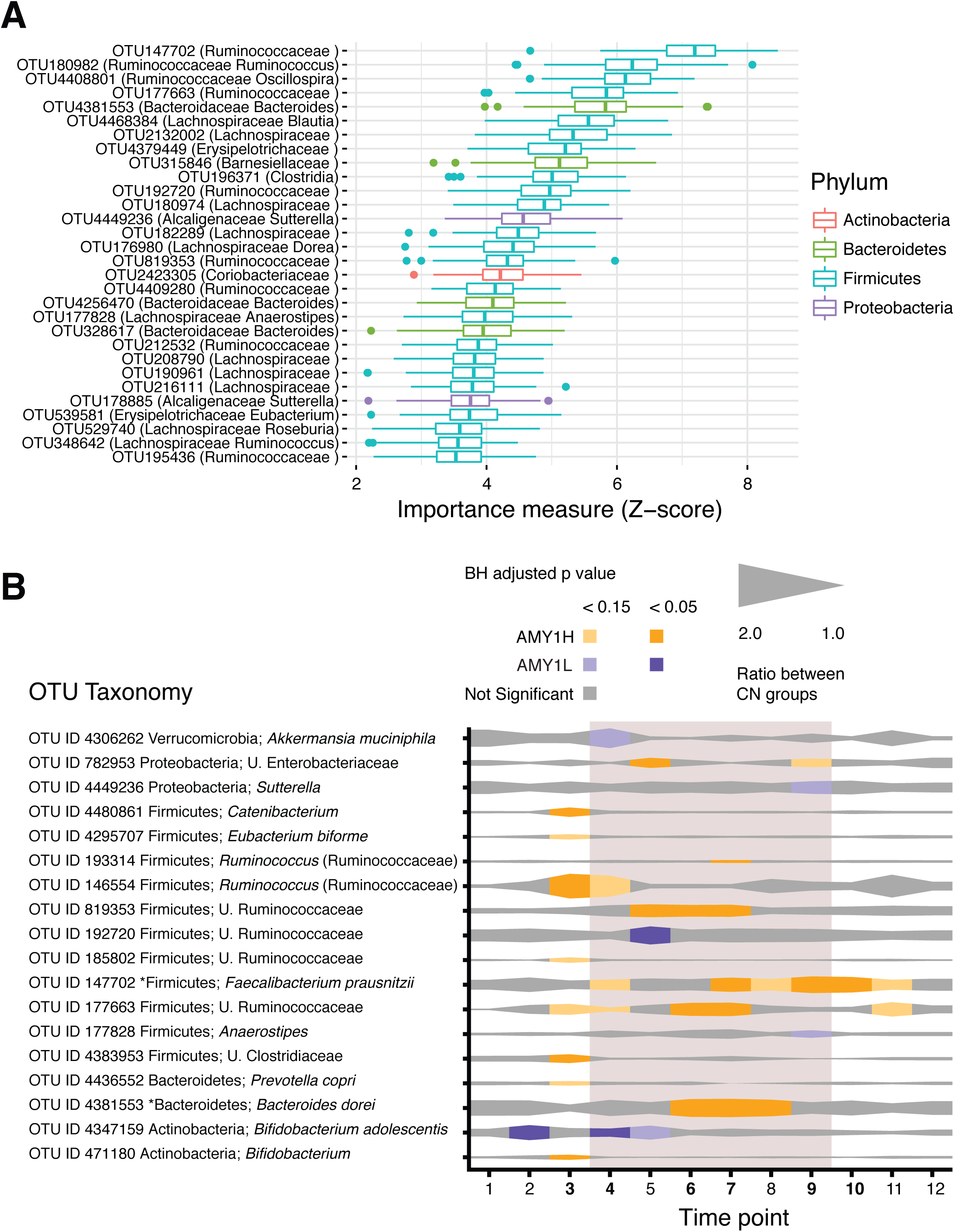
Gut microbiomes differ between SAA and *AMY1* groups at the OTU level. See legend for Figure 3. (B) Additionally, samples collected at the time points in bold print were also subjected to shotgun metagenomics analysis. See also Tables S4 and S5.

### Effect of diet on the oral and fecal microbiomes

We assessed whether diet standardization resulted in more similar microbiomes (i.e., reduced beta-diversity) between oral and fecal AMY1H and AMY1L groups by comparing samples before, during, and after the standardized diet. Non-parametric bootstrap confidence intervals (CIs) for the differences in the weighted and unweighted UniFrac distances between AMY1H and AMY1L groups for each pair of diet periods indicated that diet standardization did not induce convergence of oral microbiomes between AMY1H and AMY1L subjects, but did so for the gut microbiomes (Figure S5B,C).

### Taxa discriminatory for low/high AMY1 gut microbiomes are validated in a separate larger population

We confirmed the enrichment of discriminatory taxa in fecal microbiomes of AMY1H and AMY1L subjects British population for which host genotype (SNPs) and fecal 16S rRNA gene sequence data were available (Goodrich et al. 2016; Goodrich et al. 2014). The genotype data included 7 of the 10 SNPs that correlate with *AMY1*-CN (Usher et al. 2015). For each of the 994 British subjects with normal BMIs and available fecal 16S rRNA gene sequence data, we calculated the sum of the change in *AMY1*-CN values corresponding to each of their 7 alleles. We then selected only the top and bottom 5% of the resulting distribution to include 100 subjects: 50 with the highest and 50 with the lowest predicted total difference in *AMY1*-CN. Using Harvest, we identified 17 OTUs with significantly different relative abundances between the high and low groups (Table S5). Many of these OTUs belong to the same taxa as those enriched in gut microbiomes of AMY1H subjects (e.g., *Ruminococcus Faecalibacterium prausnitzii*, *Bacteroides*). Overall, in agreement with results from the Cornell population, members of the Ruminococcaceae family were prominent among the taxa enriched in fecal microbiomes obtained from British subjects with predicted high *AMY1*-CNs.

### Deep metagenome sequencing reveals differences in functional capacity between *AMY1*-CN gut microbiomes

We compared the metabolic potentials of the gut microbiomes for AMY1H and AMY1L groups with comparisons of fecal metagenomes generated for all subjects sampled at 6 TPs (3,4,6,7,9,10). We sub-sampled 20 million pair-end reads per sample to normalize sequencing depth and used the HMP Unified Metabolic Analysis Network (HUMAnN2) pipeline to classify shotgun metagenomic reads into gene families (Figure 6A).

**Figure 6.**
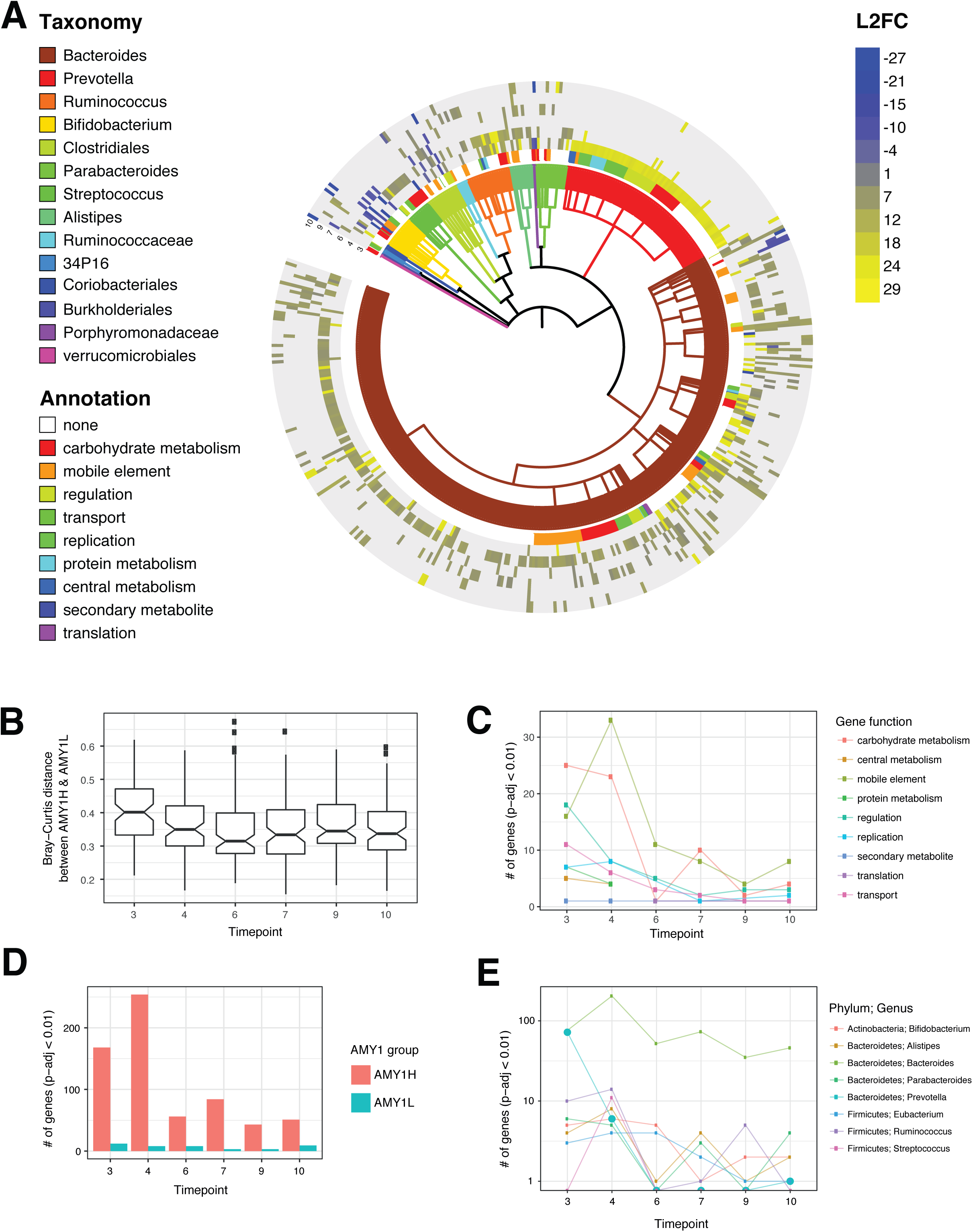
Metagenomes indicate convergence at the gene-level. (A) Heat map displaying each of 481 gene families with abundances differing between AMY1H and AMY1L at one or more of six different TPs. The heat map is sorted by taxonomy, annotation group, and gene family. Each concentric circle in the heat map corresponds to a TP. Gene family abundances with significant differences between AMY1 groups were identified using DESeq2, and the log2 fold difference between AMY1H relative to AMY1L is depicted in the heat map. Higher abundances of gene families in the AMY1H group are colored yellow, while those higher in AMY1L are colored blue. Only gene families with a BH-adjusted p < 0.01 and that were assigned taxonomy are shown. (B) Bray-Curtis distance between AMY1H and AMY1L metagenome samples (reads mapped to gene families; no taxonomic designation). (C) Number of significantly enriched gene families at each time point. (D) Number of significantly enriched gene families that could be grouped by function. (E) Taxonomy of significantly enriched gene families.

Several observations indicated that the functions of gut metagenomes converged during the period of diet provision. Using non-parametric bootstrapping with 1000 permutations, we determined that Bray-Curtis distances calculated from gene family counts decreased in mean values between AMY1H and AMY1L individuals during the diet period relative to the pre-diet period (Figure 6B). The number of gene families significantly enriched in the AMY1H group was also lower after TP 4 (Figure 6C), and the number of gene families significantly different between AMY1 groups was lower after TP 4 regardless of taxonomy or function (Figure 6D, 6E). Interestingly, we also observed a spike of differentially abundant genes associated with mobile elements at the start of the diet provision period (TP3 to TP4), which is consistent with nutritional stress-induced activation of prophages, lytic bacteriophages, and horizontal gene transfer (Huddleston 2014; Lerner et al. 2017). Thus, the diet standardization appears to have engendered a convergence of metabolic functions, consistent with the observed convergence of the gut microbial communities.

Despite the convergence of AMY1H and AMY1L microbiomes over the standardized diet period, the two groups could nevertheless be differentiated by their functional gene content. We used the statistical software DESeq2 to identify gene families with differential abundances between high and low AMY1 groups at each time point. We identified 481 gene families with significantly different read counts at one or more TPs between AMY1H and AMY1L groups with a BH-adjusted p < 0.01 (Figure 6A). Notably, 39% of the 481 gene families were taxonomically assigned to *Bacteroides dorei* and were more abundant in the AMY1H group at multiple TPs, in accordance with our 16S rRNA gene diversity results (Figures 5B, 6A). Only *B. cellulosilyticus* was enriched in the AMY1L group. In accordance with the results of the 16S rRNA gene diversity analysis, we also noted an enrichment in read counts for gene families mapping to *Ruminococcus* in the AMY1H group (Figures 5B, 6A). Sequences mapping to gene families from *P. copri* were also more abundant in AMY1H, but almost exclusively at TP 3, consistent with the 16S rRNA gene diversity data. Together, these data support a relative enrichment in taxa responsible for resistant starch breakdown in the AMY1H compared to the AMY1L gut microbiomes.

To directly assess functional capacity for carbohydrate degradation, we used hidden Markov models from dbCAN to identify Carbohydrate-Active enZYmes (CAZymes), which include the following enzyme classes: glycoside hydrolases (GH), glycosyltransferases (GT), polysaccharide lyases (PL), carbohydrate esterases (CE), carbohydrate-binding modules (CBM), S-layer homology modules of the cellulosome (SLH), and auxiliary activities (AA) (Lombard et al. 2014). We then used a linear mixed model to assess differences in the abundances of each of the 7 CAZYme classes between AMY1H and AMY1L groups. We observed a higher number of read counts for GH and PL classes in AMY1L individuals (post-hoc analysis, GH: p = 0.0054; PL: p = 0.030, Figure 7A, B), and a similar trend for AA and CE classes (Figure S6). These enzyme classes are involved in the breakdown of complex carbohydrates; their enrichment in AMY1L individuals is consistent with the notion that a greater load of complex carbohydrates reaches the distal gut in these individuals.

**Figure 7.**
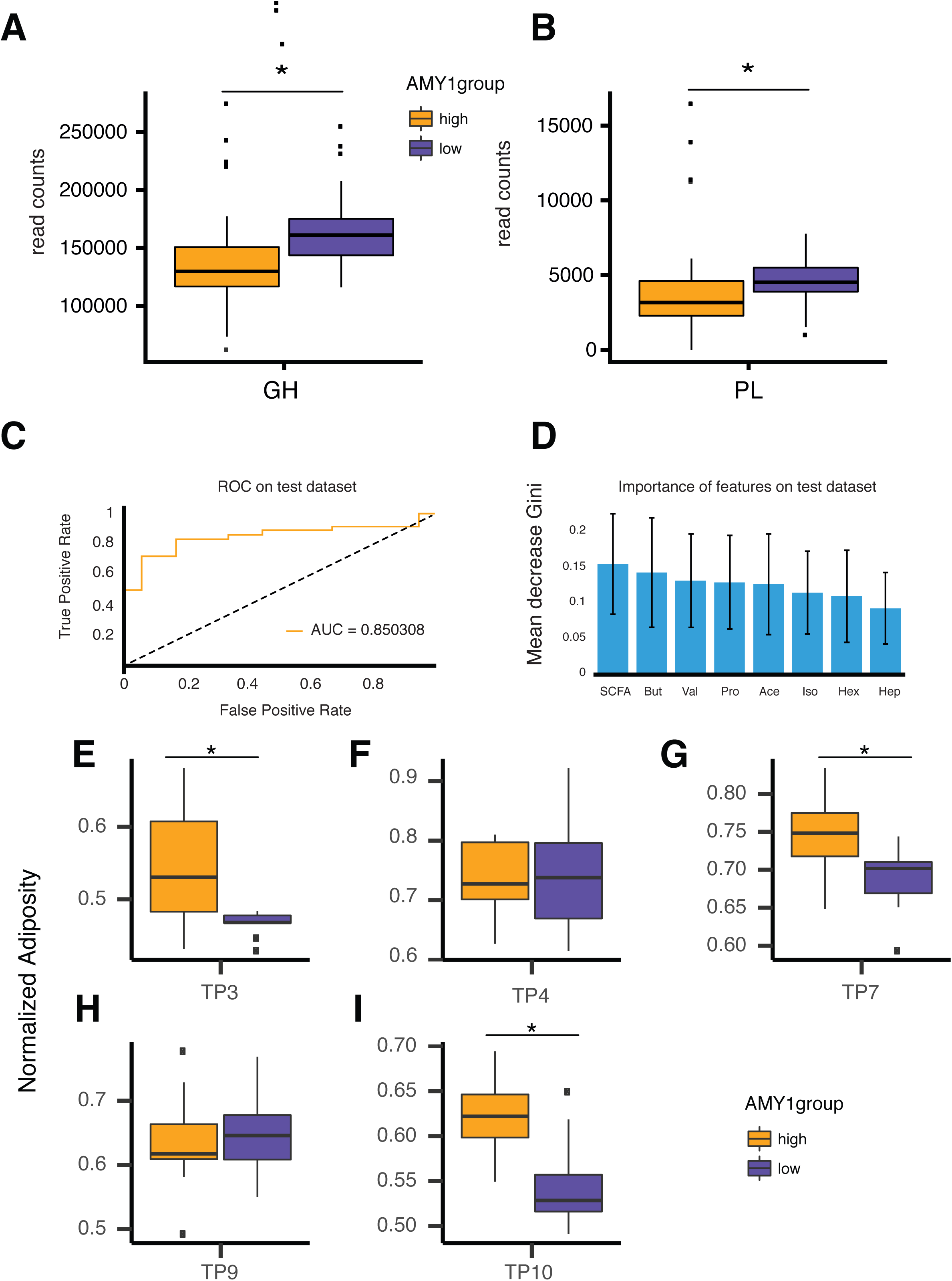
AMY1H and AMY1L gut microbiomes differ in function. (A) and (B) show boxplots of the read counts in the AMY1H and AMY1L groups for each of the significantly different Carbohydrate-Active enZYme classes: (GH) glycoside hydrolases and (PL) polysaccharide lyases. (C) We used machine learning to assess whether SCFAs levels were predictive of SAA. Subjects were clustered according to their salivary amylase activity in two groups (SAA-H and SAA-L) by k-means. Shown here is the receiver operating characteristic (ROC) curve of a random forest model used to predict the SAA group using the SCFAs measurements in a test dataset. (D) The features used by the random forest model for the classification of the test samples are shown in decreasing order of importance given by the mean decrease in Gini index. SCFA is the total of all SCFAs. But = butyrate, Val = valerate, Pro = propionate, Ace = acetate, Iso = isovalerate, Hex = hexanoate, Hep = heptanoate. (E) - (I) Boxplots of the adiposity measure normalized by baseline weight on the day of inoculation of formerly germfree mice after humanization with stool collected from the study participants at 5 time points. Time point is indicated on the x axis. * Tukey’s HSD adjusted p < 0.05.

### Fecal short chain fatty acids relate to salivary amylase activity

As an assessment of microbial metabolic output, we measured levels of short chain fatty acids (SCFAs) in stool samples collected at all TPs. We used machine learning to assess whether SCFAs levels were predictive of SAA (high and low SAA determined by k-means clustering for all 25 subjects; all observations from the same subject were labeled with the subject’s SAA group). By using SAA groups instead of AMY1 groups, we correlated data measurements that were collected over time and exhibited day-to-day variation. We trained a random forest model with 80% of the available SCFA measures to predict the SAA group to which the remaining 20% belonged. The model achieved an accuracy of 83.61% with a MCC of 64.46%, and an AUC of 85.03% (Figure 7C). The total concentration of SCFAs was the most informative element for discriminating the high and the low SAA groups, followed by the concentrations of butyrate, valerate, propionate, and acetate (Figure 7D). Given that *AMY1*-CN is positively correlated to SAA, we ran the same models using the *AMY1*-CN instead of SAA: results were not significant although they showed a similar trend (data not shown). Using a linear mixed model that included SAA group as a covariate, we confirmed that the concentrations of several SCFAs were higher in subjects with high SAA (adjusted p-values: total SCFA concentration = 4.7×0^−2^, acetate = 6.5×0^−2^, propionate = 6.5×0^−2^, butyrate = 4.7×0^−2^). Assuming equal uptake of SCFAs in the colon across subjects, these results suggest that the higher the host SAA, the greater the SCFA production in the colon. Given that SAA can vary from day to day for a given individual, the observation that SAA is a better predictor of SCFAs than *AMY1*-CN indicates that the microbiome’s metabolite output is sensitive to daily SAA variation.

### Fecal transplants from AMY1H donors into germfree mice promote greater adiposity

To gauge differences in function for gut microbiomes at the high and low ends of the *AMY1*-CN distribution, we used fecal transplantation from AMY1H and AMY1L donors, sampled at 5 TPs, to 96 male Swiss-Webster 4-6 week old germfree mice fed a polysaccharide-rich chow *ad-libitum* and single-caged post-transfer. Adiposity was assessed by DEXA after 4-6 weeks. Across all mice, we observed a significantly higher body fat percentage for recipients of the AMY1H compared to the AMY1L microbiomes (linear mixed model; p = 0.026). Post hoc pairwise comparisons revealed, that TPs 3, 7 and 10 showed a significantly higher final adiposity for the AMY1H compared to the AMY1L treatment groups (Tukey’s HSD adjusted p < 0.05), whereas TPs 4 and 9 did not (after controlling for weight on the day of inoculation and length of the experiment; Figures 7E-I). Food intake was not significantly different between high versus low *AMY1*-CN donor groups and there were no differences in intestinal inflammation (measured by Lipocalin 2 at the end of the experiments using samples from TPs 3 and 10, data not shown). Thus, the AMY1H microbiomes generally drove higher adiposity gains that were unrelated to food intake and metabolic inflammation.

## Discussion

Our study shows that variation in the copy number of the human salivary amylase gene *AMY1* is associated with the diversity and function of the human oral and gut microbiome. The *AMY1*-CN of the host directly influences the carbohydrate milieu in the GI tract through its dose-dependent effect on salivary amylase production. We show here that *AMY1*-CN is associated with the composition of the oral microbiome and with the composition and function of the gut microbiome.

Host AMY1-CN impacted the oral microbiome in a diet-independent manner. Our month-long study included a 2-week period in which the subjects were supplied with all meals, and we asked them to record any other food consumed. This diet intervention did not regulate the amount of food consumed, and did not preclude consumption of unreported food, but it did ensure that subjects consumed carbohydrate-rich foods daily. The oral microbiomes, first to experience food intake, were unaffected by the standardized diet period. They were, however, sensitive to host *AMY1*-CN status and to SAA. Notably, compared to AMY1L, AMY1H subjects harbored higher levels of *Porphyromonas sp.* (e.g., *P. endodontalis*). Several members of the genus *Porphyromonas,* including *P. endodontalis*, have been associated with periodontitis (Socransky et al. 1998; Park et al. 2015; Colombo et al. 2012; Wade 2013; Griffen et al. 2012; Lombardo Bedran et al. 2012; Cao et al. 2012). Many of the same taxa were discriminatory for host SAA, across all subjects and TPs assayed. Interestingly, we did not detect higher relative abundances of OTUs mapping to the genus *Streptococcus* in the AMY1H group, possibly because the majority of the *Streptococcus* OTUs were unclassifiable at the species level (an issue that has been previously reported even with full-length 16S rRNA gene sequences (Wade 2013; Hanage, Fraser, and Spratt 2005). The strong relationship of the oral microbiota composition to host *AMY1*-CN and SAA, coupled to the insensitivity to the change in diet, indicates a long-term conditioning of oral microbiomes to their host’s average SAA, which is regulated by the *AMY1*-CN.

In contrast to the oral microbiomes, the standardized diet period had a noticeable effect on the composition of the gut microbiomes. The convergence was most dramatic in the functional profiles of the microbiomes, and less detectable at the OTU level. Differences in the strength of this signal can be attributed to differences in resolution at the species and functional levels (Noecker et al. 2017). Convergence of the gut microbiomes of hosts consuming similar diets has previously been observed (Muegge et al. 2011; Clayton et al. 2016; Minot et al. 2011). In our study, the onset of the diet was accompanied by a spike in the mobile-gene elements of the fecal metagenomes, similar to what has been described for stress responses (Huddleston 2014; Lerner et al. 2017).

Despite the convergence of community and function on the gut microbiomes, the *AMY1*-CN status of the host was reflected in its composition and function. For instance, members of the Ruminococcaceae family, which along with some *Bacteroides spp* have been reported to ferment resistant starch (Walker et al. 2011; Salonen et al. 2014; Herrmann et al. 2017; Moraïs et al. 2016; Ze et al. 2012; Flint et al. 2012; Upadhyaya et al. 2016), were enriched in the AMY1H microbiomes. We validated these results in a separate and much larger population of British subjects for whom *AMY1*-CN was estimated from predictive SNPs, and 16S rRNA data was also generated by our laboratory using the same protocols (Goodrich et al., 2016). Many of the same taxa were enriched in the subjects at the high end compared to the low end of the predicted *AMY1*-CN gradient formed by the British population. Thus, association of taxa involved in the degradation of resistant starch in host with high *AMY1*-CN may be a widespread feature of gut microbiomes.

Host *AMY1*-CN was also related to the functional capacity of the gut microbiome. Our metagenomic analysis revealed two classes of carbohydrate-active enzymes involved in the breakdown of complex carbohydrates in general, glycoside hydrolases and polysaccharide lyases, as enriched in the microbiomes of AMY1L hosts. This observation is reminiscent of what has been noted in urbanized versus rural populations (Mancabelli et al. 2017). These results suggest that for a given diet, the AMY1L distal gut microbiota may be presented with a greater load of complex carbohydrates than the AMY1H microbiota. In contrast, as a result of greater average host SAA, the proportion of resistant starch is greater in the AMY1H colon, and the corresponding fermenters (e.g., *Ruminococcus* and *Bacteroides*) are more abundant.

We assessed functional output of AMY1H and AMY1L microbiomes with measures of SCFAs. SCFAs are fermentation products of distal gut microbiota, and their levels in stool are influenced both by production by the microbiome and uptake by the host. We observed that SCFAs in stool were associated more strongly with host SAA levels than with host *AMY1*-CN. Within an individual, SAA varies from day to day. Although *AMY1*-CN is a good predictor of average SAA, it may be a poor predictor of SAA on any given day. A better association of SCFAs with SAA compared to *AMY1*-CN indicates that fecal SCFA pools reflect short-term fermentation dynamics in the gut that are affected by fluctuating SAA. Since microbiota known to ferment resistant starch (e.g., Ruminococcaceae) are enriched in the AMY1H subjects, the activity of these microbiota may be driving the higher levels of SCFAs in their stool (Topping and Clifton 2001).

Another way we tested the functional capacity of the gut microbiomes was to transfer fecal microbiota to germfree mice and assess adiposity differences for mice receiving microbiomes of AMY1H compared to those receiving AMY1L microbiomes. We transplanted human microbiomes 1:1 into mice, and used 5 samples per subject. The 5 transfer experiments are not exact replicates, because we used five separate samples collected at different TPs from each donor, which takes into account daily variability in microbiomes. Overall, we observed a greater adiposity for mice recipients of microbiomes derived from the AMY1H donors. When results for each TP are considered separately, three out of five TPs are driving this effect. The overall significant difference in mean adiposity between treatment groups provides evidence that AMY1H microbiomes transplanted into mice more efficiently extract calories from the mouse diet.

Given that the AMY1L microbiomes were enriched in taxa and gene functions associated with complex carbohydrate breakdown, it may seem surprising that the AMY1H microbiome transfers led to higher average adiposity in germfree mouse recipients. The mice used here are AMY1L (CN = 2) and consumed a diet rich in complex carbohydrates, including cellulose. The AMY1H donor microbiomes featured higher levels of *Ruminococcus spp.*, some of which have been reported to degrade cellulose in the human gut (Chassard et al. 2010). Thus, one possible explanation is that the human AMY1H-conditioned microbiomes more efficiently degraded the cellulose and other insoluble fiber contained in the mouse diet. The mouse transfers provide a functional read-out by allowing strict control of *AMY1*-CN and diet, and results highlight how a compositional difference in microbiomes can translate to differences in functional output. Within their native human hosts, however, AMY1H microbiomes may not behave the same way (i.e., more obesogenic than AMY1L), since they are not decoupled from their high SAA environment.

### Prospectus

Our results may be particularly pertinent to current efforts to tailor diet to individual microbiomes. One of the strongest drivers of gut microbial community composition is dietary intake (e.g. David et al. 2014; Wu et al. 2011; Duncan et al. 2007; Muegge et al. 2011). Some microbes have been shown to preferentially degrade specific forms of polysaccharide (Flint et al. 2012; Cockburn and Koropatkin 2016; Walker et al. 2011). Several of these polysaccharides, including resistant starch, affect host physiological parameters including glucose homeostasis and adiposity (Bindels et al. 2015; Keenan et al. 2015; Birt et al. 2013). Traditionally, the glycemic index of a given food has been reported as a single value assumed to be the same for all individuals, with the intent of identifying carbohydrates with a lower glycemic index as being healthier. However, host glucose and gut microbial responses to various starches have exhibited marked interindividual variation between human subjects (Walker et al. 2011; Martínez et al. 2010; Venkataraman et al. 2016; Korem et al. 2017; Kovatcheva-Datchary et al. 2015). Host genetic factors, particularly *AMY1*-CN, could be contributing to these differences in glucose response and microbiome composition during controlled dietary interventions. Future studies aimed at choosing carbohydrates types and amounts based on the gut microbiome may seek to incorporate host *AMY1*-CN as an important interaction variable.

Finally we note that selection for duplication at the *AMY1* locus and lactase persistence evolved around the Neolithic Transition to an agrarian lifestyle approximately 10,000 years ago. Recent genome-wide association studies have linked allelic variation to microbiome composition, and the strongest of these associations is between the *LCT* gene (encoding lactase) and levels of the lactose-degrading *Bifidobacterium* genus in the gut (Blekhman et al. 2015; Goodrich et al. 2016; Bonder et al. 2016). This study, together with the studies linking lactase gene alleles to levels of Bifidobacteria, underscore how recent human adaptation to new diets drove human genetic variation across populations and underlies differences in modern-day microbiomes.

### Author Contributions

R.E.L. supervised the study; R.E.L. and A.C.P. designed the study and oversaw sample collection. A.C.P., J.L.S., and Q.S. generated data; A.C.P. performed the statistical analyses with contributions from J.K.G., L.M.J., N.D.Y., A.R., G.G.L., H.Y.B., M.E., D.H.H., R.E.L., and J.G.B. A.C.P., S.L., J.L.W., J.K.G, and Q.S. performed microbiota transfer experiments. A.C.P., J.K.G., N.D.Y., A.R., and G.G.L. and R.E.L. prepared the manuscript.

## Acknowledgements

We thank Brenda Daniels-Tibke and Kacie Harrington of the Cornell’s Human Metabolic Research Unit, and Fiona Hoi Yi Chan for assistance with the standardized diet data. We thank Sha Li, Noah Clark, M. Elizabeth Bell, Wei Zhang, Tim DeMarsh, and Julia Hildebrandt for technical assistance. This work utilized Core Services supported by grant DK097153 of NIH to the University of Michigan. This work was supported by a David and Lucile Packard Foundation Fellowship (R.E.L.), the Arnold and Mabel Beckman Foundation (R.E.L.), a National Science Foundation Graduate Fellowship (J.K.G.), The Hartwell Foundation (R.E.L.), and the Max Planck Society.

---

**Table S1.**
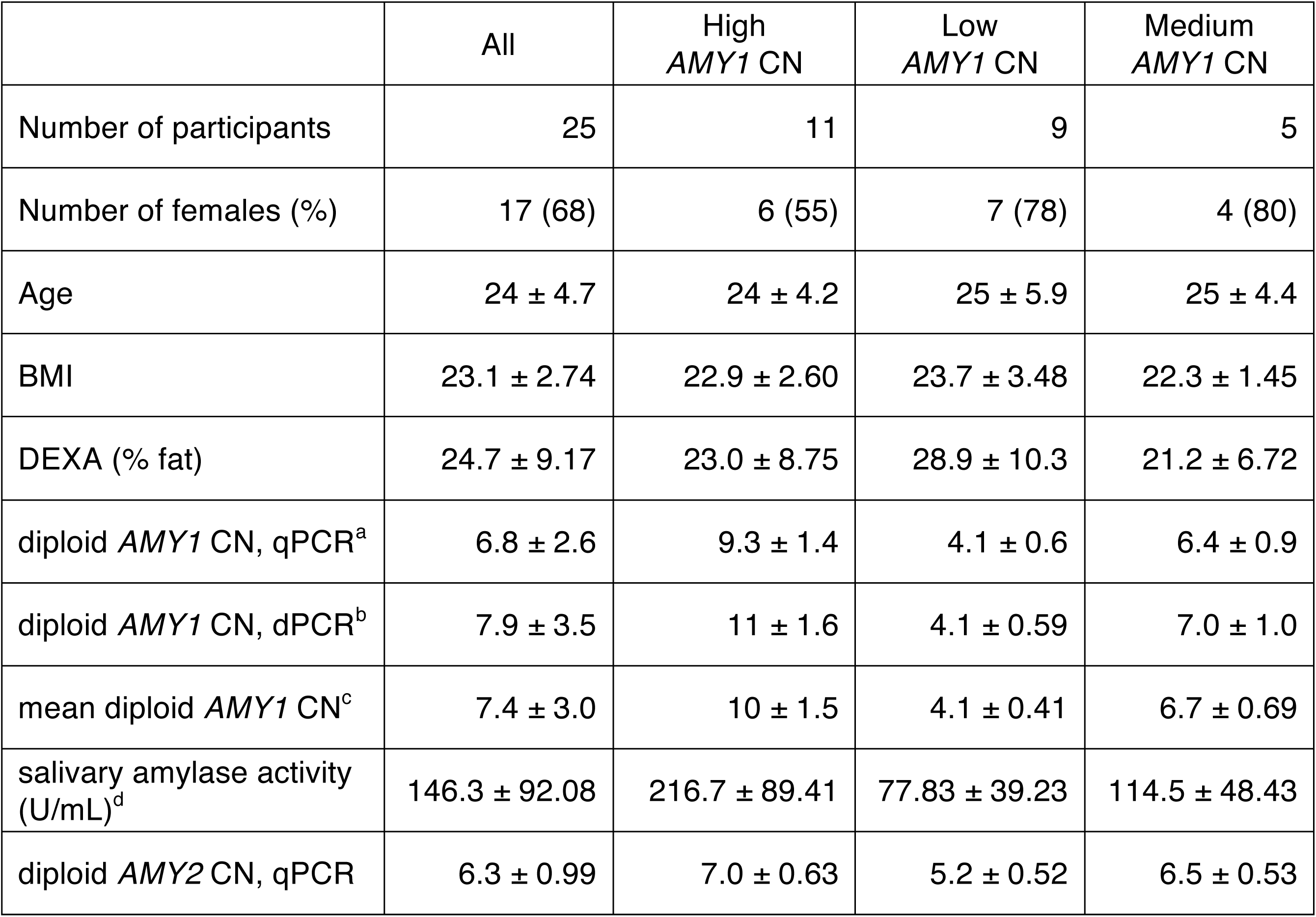
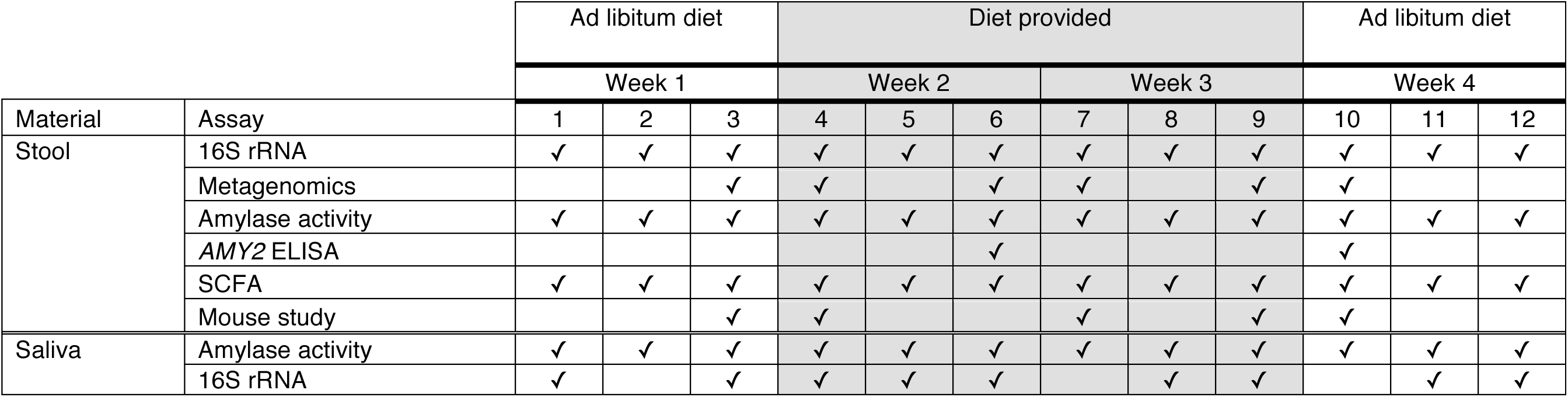
Related to Table 1. Assays performed on samples collected during the study. Stool and saliva samples were collected at 12 time points throughout the 4-week study, which included two weeks when meals and snacks were provided. Assays performed on each sample type are shown here.

**Table S2.** Related to Table 1 and Figure S1. Menus detailing the diet provided to participants during week 2 and week 3 of the study. Meals and snacks provided during the 2-week diet provision period. Days 1 through 7 were repeated during the second week.

|  | **Breakfast** | **Lunch** | **Dinner** |
| --- | --- | --- | --- |
| **Day 1** | • Blueberry pancakes<br>• Butter<br>• Maple syrup<br>• Milk | • Ham & Swiss cheese sandwich on multi-grain bread (Vegetarian option: Seitan & Swiss cheese sandwich on multi-grain bread)<br>• Carrot sticks<br>• Milk | • Spaghetti with ground turkey and marina sauce (Vegetarian option: Quinoa stuffing)<br>• Grated parmesan cheese |
| **Day 2** | • Whole grain toast<br>• Jam<br>• Hard-boiled eggs | • Hummus & veggie wrap<br>• Apple | • Soy-ginger beef stir-fry (Vegetarian option: Tofu stir-fry with rice, broccoli, peppers and onions) |
| **Day 3** | • Instant oatmeal with banana, raisins & walnuts | • Turkey, pesto and Provolone cheese sandwich (Vegetarian option: Provolone cheese and pesto sandwich w/ cucumber) | • 2 Tacos (Vegetarian option: 2 Bean-Vegetable Tacos)<br>• Salsa<br>• Sour cream |
| **Day 4** | • Bowl of Go Lean Crunch cereal<br>• Low-fat milk<br>• Orange | • Chicken salad sandwich (Vegetarian option: Asian Kasha salad over mixed greens)<br>• Grapes | • Chicken cacciatore over bow tie pasta (Vegetarian option: Mixed-bean cacciatore over bow tie pasta) |
| **Day 5** | • Vanilla Greek yogurt<br>• Cantaloupe melon<br>• Granola | • Black bean burrito with rice and corn and cheese<br>• Salsa | • Greek salad with sautéed chicken (Vegetarian option: Greek salad with taboule and hummus) |
| **Day 6** | • Bagel<br>• Peanut butter<br>• Banana<br>• Milk | • Egg salad on pita bread<br>• Carrots | • Baked chicken (Vegetarian option: Baked tofu)<br>• String beans<br>• Baked sweet potato |
| **Day 7** | • Eggs<br>• Whole wheat<br>• English muffin<br>• Butter<br>• Jam | • Turkey, apple, and cranberry chutney sandwich (Vegetarian option: Brie, apple, and cranberry chutney sandwich) | • Veggie pizza on whole wheat crust |

Snacks (every day options): String cheese, Oranges, Apples, Nature valley granola bars, Juice, and Pretzels

**Table S3.**
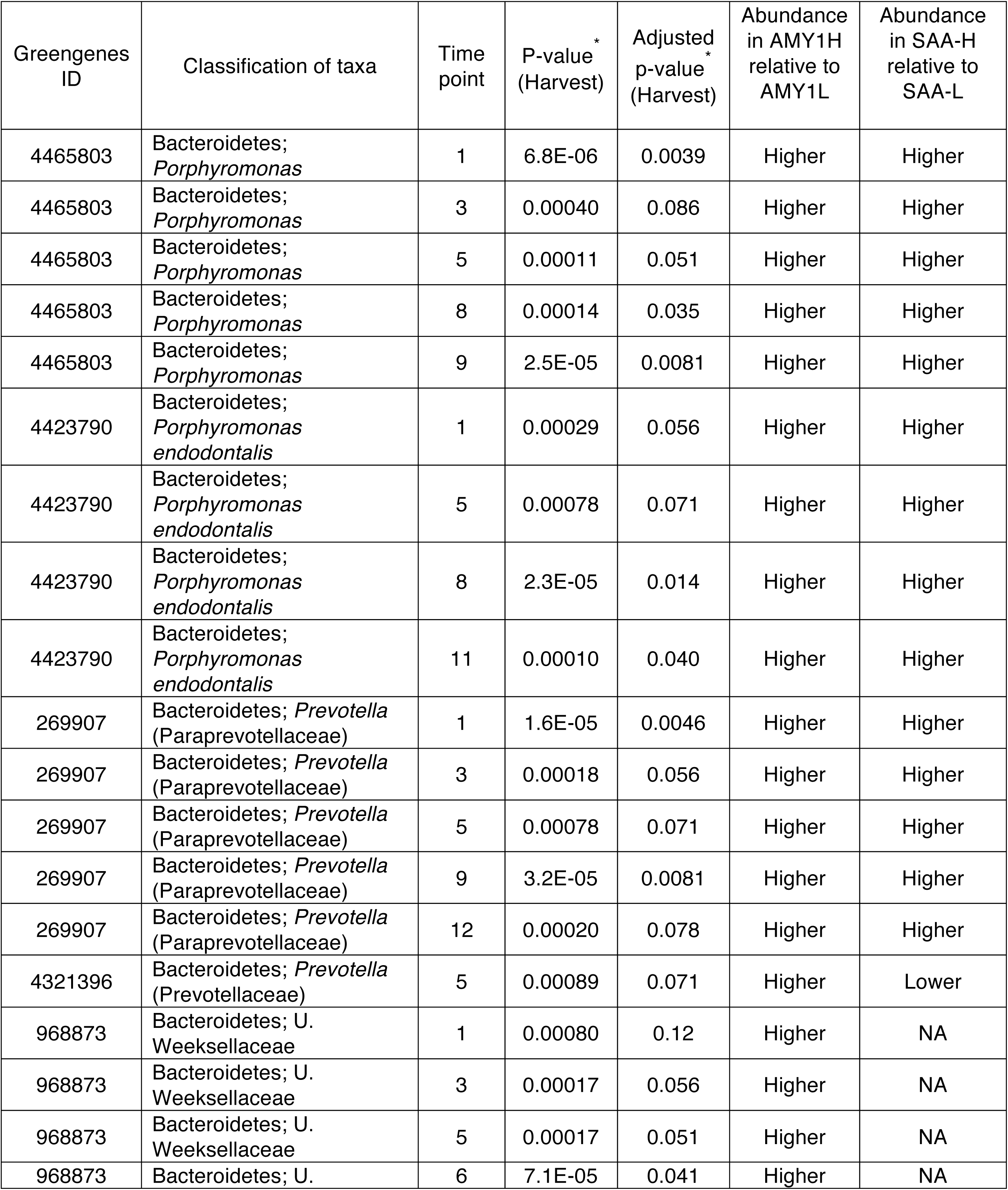
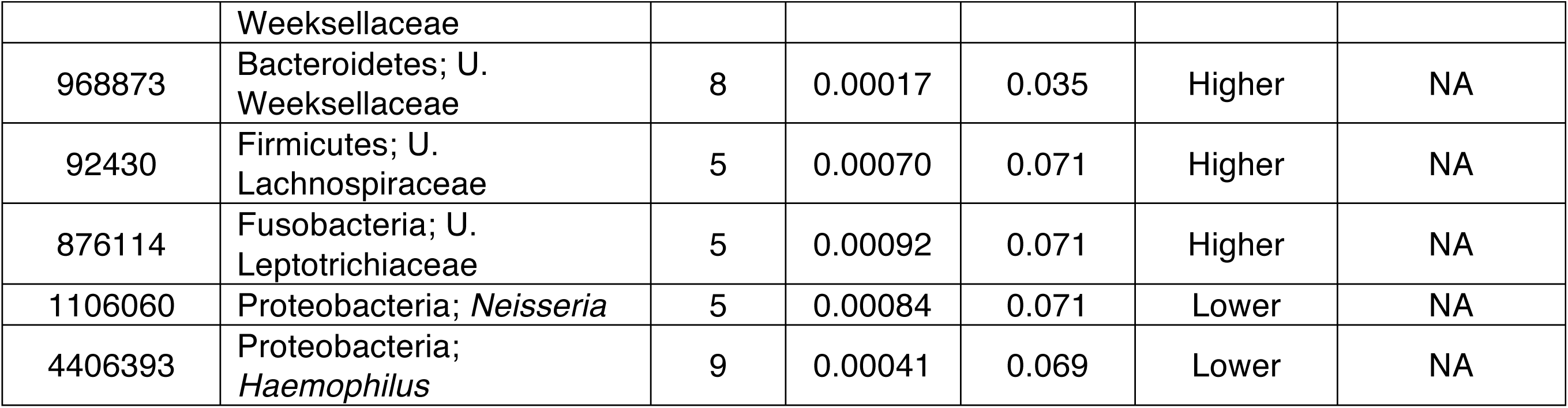
Related to Figure 3. Differentially abundant taxa in the saliva of Cornell participants. Greengenes ID numbers for the OTUs with differential abundances at one or more time points identified using Harvest. The AMY1 group in which each OTU was enriched is indicated as well as the SAA group of enrichment for OTUs also identified in the machine learning analysis. *The nominal and BH-adjusted p values obtained using Harvest are shown.

**Table S4.** Related to Figure 5. Differentially abundant taxa in the stool of Cornell participants. See legend for Table S3.

| Greengenes ID | Classification of taxa | Time point | P-value<sup>*</sup><br>(Harvest) | Adjusted p-value<sup>*</sup><br>(Harvest) | Abundance in AMY1H relative to AMY1L | Abundance in SAA-H relative to SAA-L |
| --- | --- | --- | --- | --- | --- | --- |
| 146554 | Firmicutes; *Ruminococcus* | 3 | 9.4E-06 | 0.0050 | Higher | NA |
| 146554 | Firmicutes; *Ruminococcus* | 4 | 7.8E-05 | 0.051 | Higher | NA |
| 147702 | Firmicutes; Unclassified Ruminococcaceae | 2 | 0.00040 | 0.16 | Higher | Higher |
| 147702 | Firmicutes; Unclassified Ruminococcaceae | 4 | 0.00048 | 0.13 | Higher | Higher |
| 147702 | Firmicutes; Unclassified Ruminococcaceae | 7 | 9.7E-05 | 0.029 | Higher | Higher |
| 147702 | Firmicutes; Unclassified Ruminococcaceae | 8 | 0.00023 | 0.14 | Higher | Higher |
| 147702 | Firmicutes; Unclassified Ruminococcaceae | 9 | 3.1E-05 | 0.040 | Higher | Higher |
| 147702 | Firmicutes; Unclassified Ruminococcaceae | 10 | 1.2E-05 | 0.013 | Higher | Higher |
| 147702 | Firmicutes; Unclassified Ruminococcaceae | 11 | 0.00022 | 0.11 | Higher | Higher |
| 177663 | Firmicutes; Unclassified Ruminococcaceae | 3 | 0.00068 | 0.12 | Higher | Higher |
| 177663 | Firmicutes; Unclassified Ruminococcaceae | 4 | 0.00035 | 0.12 | Higher | Higher |
| 177663 | Firmicutes; Unclassified Ruminococcaceae | 6 | 7.7E-06 | 0.0082 | Higher | Higher |
| 177663 | Firmicutes; Unclassified Ruminococcaceae | 7 | 8.9E-06 | 0.0070 | Higher | Higher |
| 177663 | Firmicutes; Unclassified Ruminococcaceae | 11 | 0.00025 | 0.11 | Higher | Higher |
| 177828 | Firmicutes; *Anaerostipes* | 9 | 0.00031 | 0.12 | Lower | NA |
| 185802 | Firmicutes; Unclassified Ruminococcaceae | 3 | 0.0011 | 0.14 | Higher | NA |
| 192720 | Firmicutes; Unclassified Ruminococcaceae | 5 | 2.4E-05 | 0.012 | Lower | Lower |
| 193314 | Firmicutes; *Ruminococcus* | 7 | 1.6E-05 | 0.0070 | Higher | Lower |
| 471180 | Actinobacteria; *Bifidobacterium* | 3 | 0.00018 | 0.049 | Higher | Lower |
| 782953 | Proteobacteria; Unclassified Enterobacteriaceae | 5 | 5.7E-06 | 0.0085 | Higher | Lower |
| 782953 | Proteobacteria; Unclassified Enterobacteriaceae | 9 | 0.00036 | 0.12 | Higher | Lower |
| 819353 | Firmicutes; Unclassified Ruminococcaceae | 5 | 2.1E-05 | 0.012 | Higher | Lower |
| 819353 | Firmicutes; Unclassified Ruminococcaceae | 6 | 1.3E-05 | 0.0082 | Higher | Lower |
| 819353 | Firmicutes; Unclassified Ruminococcaceae | 7 | 9.7E-06 | 0.0070 | Higher | Lower |
| 4295707 | Firmicutes; *Eubacterium biforme* | 3 | 0.00048 | 0.10 | Higher | NA |
| 4306262 | Verrucomicrobia; *Akkermansia muciniphila* | 4 | 0.00014 | 0.062 | Lower | NA |
| 4347159 | Actinobacteria; *Bifidobacterium adolescentis* | 2 | 3.2E-05 | 0.038 | Lower | Lower |
| 4347159 | Actinobacteria; *Bifidobacterium adolescentis* | 4 | 1.6E-05 | 0.021 | Lower | Lower |
| 4347159 | Actinobacteria; *Bifidobacterium adolescentis* | 5 | 0.00020 | 0.075 | Lower | Lower |
| 4381553 | Bacteroidetes; *Bacteroides* | 2 | 0.00034 | 0.16 | Higher | Higher |
| 4381553 | Bacteroidetes; *Bacteroides* | 6 | 4.3E-05 | 0.018 | Higher | Higher |
| 4381553 | Bacteroidetes; *Bacteroides* | 7 | 1.9E-05 | 0.0070 | Higher | Higher |
| 4381553 | Bacteroidetes; *Bacteroides* | 8 | 3.2E-07 | 0.00039 | Higher | Higher |
| 4383953 | Firmicutes;Unclassified Clostridiaceae | 3 | 3.4E-05 | 0.012 | Higher | Lower |
| 4436552 | Bacteroidetes; *Prevotella copri* | 3 | 0.00079 | 0.12 | Higher | Lower |
| 4449236 | Proteobacteria; *Sutterella* | 9 | 0.00011 | 0.072 | Lower | Higher |
| 4480861 | Firmicutes; *Catenibacterium* | 3 | 1.4E-06 | 0.0015 | Higher | Lower |

**Table S5.** Related to Figure 5. Greengenes ID numbers for the OTUs with differential abundances previously determined for stool samples obtained for a British population were identified using Harvest. The AMY1 group in which each OTU was enriched is indicated.

| Greengenes ID | Classification of taxa | P-value | Adjusted p-value | Abundance in AMY1H relative to AMY1L |
| --- | --- | --- | --- | --- |
| 4405146 | Firmicutes; U. Clostridiales | 4.2E-06 | 0.0042 | Higher |
| 1102370 | Firmicutes; U. Ruminococcaceae | 1.2E-05 | 0.0062 | Higher |
| 174611 | Firmicutes;<br>*Faecalibacterium prausnitzii* | 0.00010 | 0.033 | Higher |
| 173810 | Firmicutes; U. Ruminococcaceae | 0.00013 | 0.033 | Higher |
| 178845 | Firmicutes; U. Ruminococcaceae | 0.00026 | 0.053 | Higher |
| 4300690 | Firmicutes; U. Ruminococcaceae | 0.00031 | 0.053 | Higher |
| 1952 | Bacteroidetes;<br>*Parabacteroides* | 0.00038 | 0.055 | Lower |
| 193191 | Firmicutes; *Lachnospira* | 0.00049 | 0.062 | Higher |
| 193969 | Firmicutes; U. Clostridiales | 0.00072 | 0.080 | Higher |
| 4412540 | Firmicutes; U. Ruminococcaceae | 0.00080 | 0.080 | Higher |
| 4396292 | Firmicutes; U. Ruminococcaceae | 0.00010 | 0.087 | Higher |
| 181059 | Firmicutes; U. Clostridiales | 0.0012 | 0.097 | Higher |
| 199534 | Bacteroidetes; *Bacteroides* | 0.0013 | 0.098 | Higher |
| 4465124 | Firmicutes; *Clostridium* | 0.0015 | 0.098 | Higher |
| 184534 | Firmicutes; U.<br>Christensenellaceae | 0.0015 | 0.098 | Higher |
| 4336939 | Firmicutes; U.<br>Ruminococcaceae | 0.0019 | 0.12 | Higher |
| 4468466 | Firmicutes; *Oscillospira* | 0.0024 | 0.14 | Higher |

**Figure S1.**
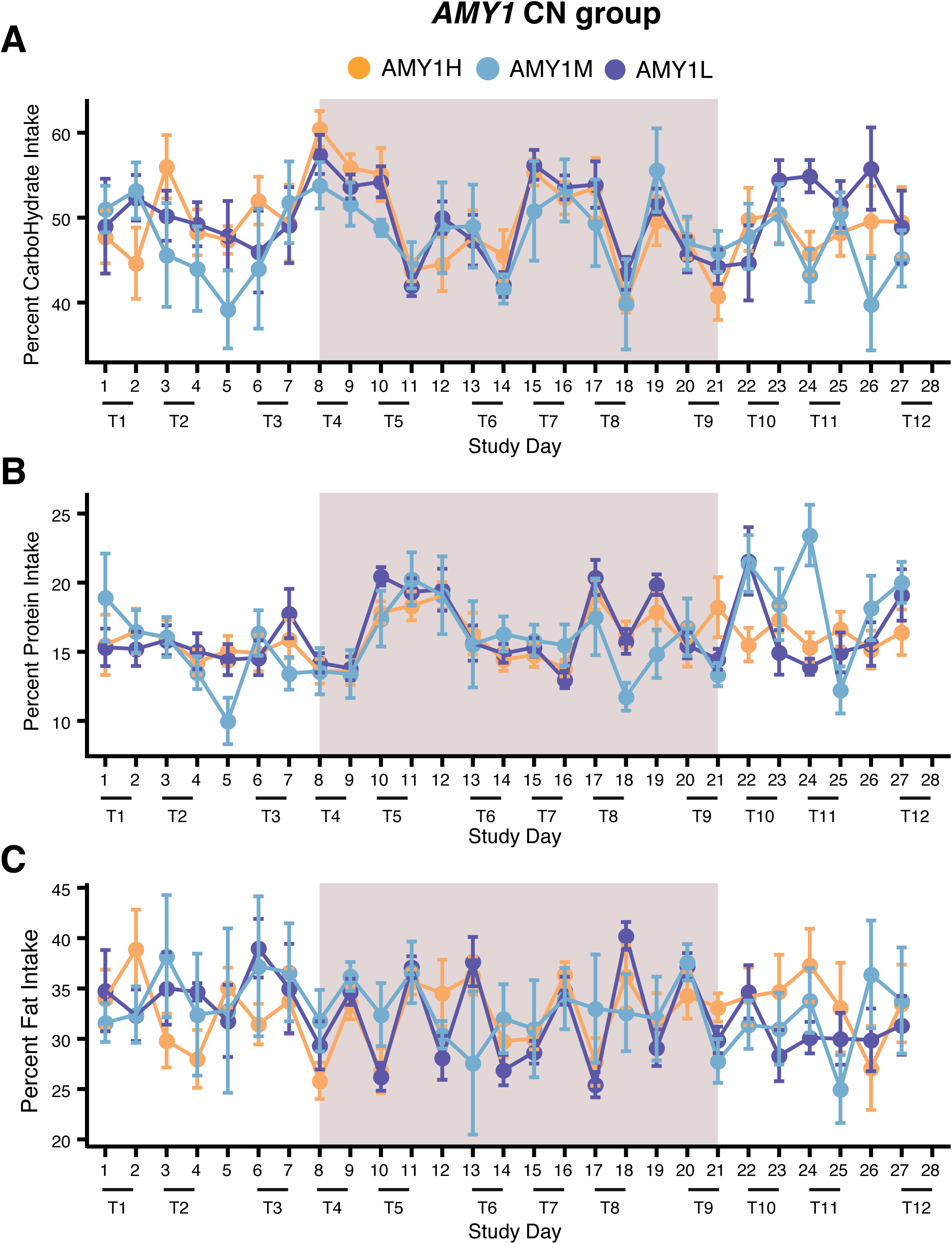
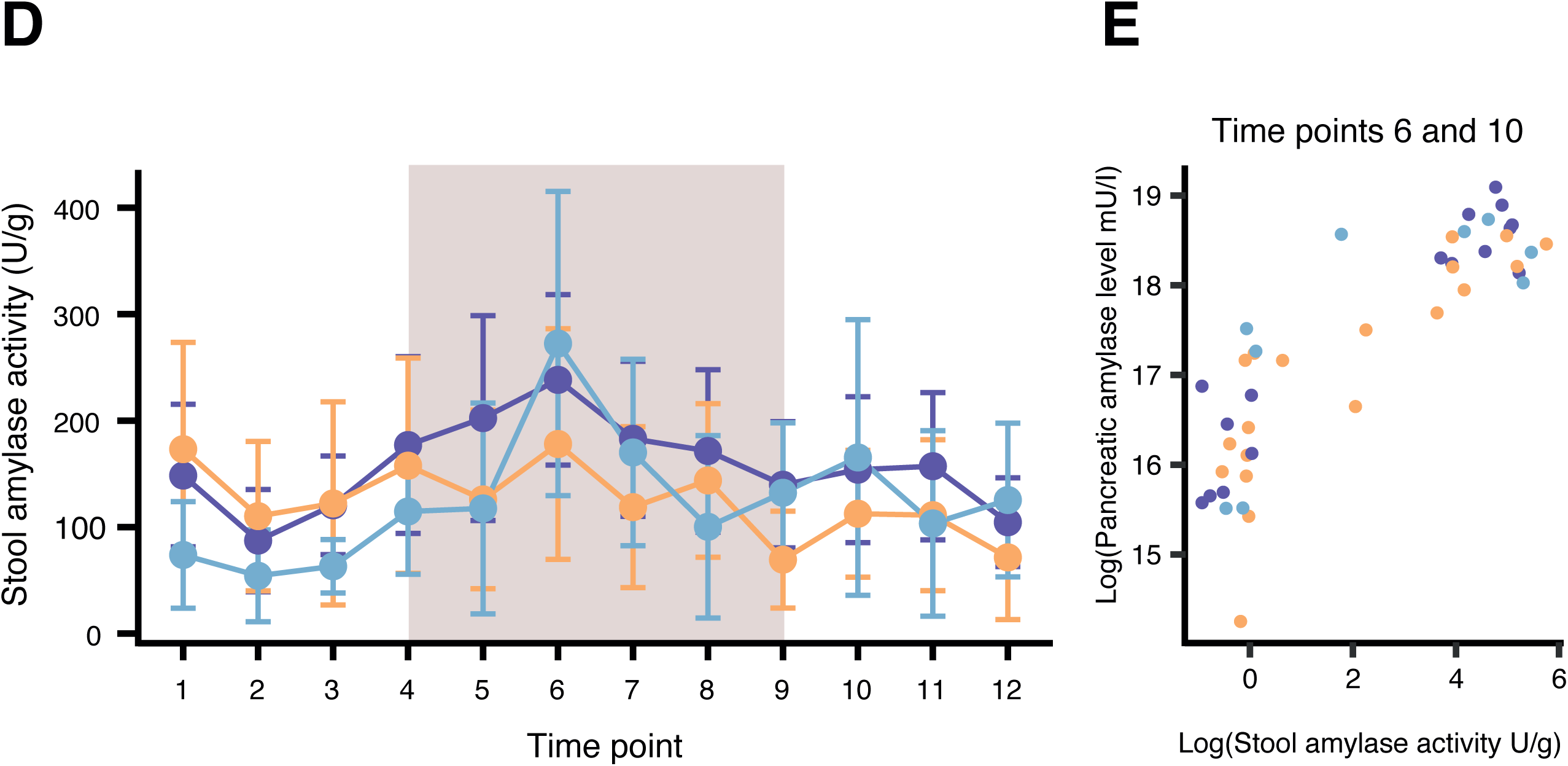
Related to Table 1. Macronutrient intake did not differ between AMY1H and AMY1L groups. Dietary intake on each day of the study for all subjects was manually entered into the nutritional analysis software SuperTracker. SuperTracker provides (A) carbohydrate, (B) protein, and (C) fat intake as a percentage of total calories. Shown here are the mean percentages ± SEM for the AMY1H, AMY1L, and medium *AMY1* CN groups. The rectangle in the background delineates the days on which diet was provided. Diet provision began with lunch on day 8 and ended with breakfast on day 22. Data for day 28 were not included in the analyses because not all subjects provided dietary intake records for the entire day. Days when stool and saliva were collected (12 TPs) are indicated at the bottom of the graph. (D) Amylase activity measurements in stool for the AMY1 groups over time (mean ± SEM). (E) Stool amylase activity versus *AMY2* enzyme levels for human stool samples collected at TPs 6 and 10 are displayed on a logarithmic scale.

**Figure S2.**
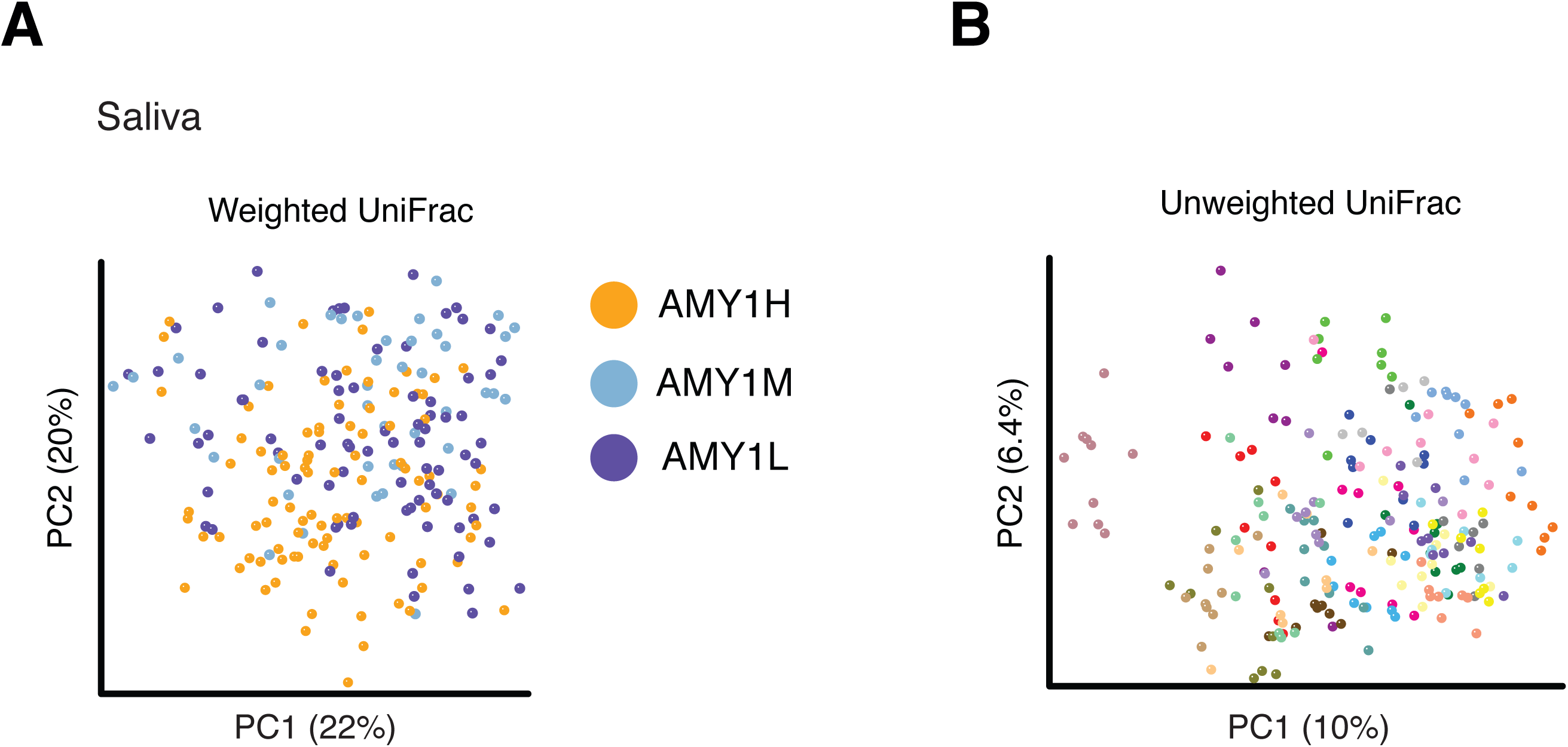
Related to Figure 2. Principal coordinates analysis of saliva samples. Principal coordinates analysis (PCoA) of the saliva samples (A) using weighted UniFrac distances and colored by AMY1 group and (B) using unweighted UniFrac distances and colored by donor, i.e. all samples from a single individual are the same color. PCs show the amount of variation explained. Panels include samples from all time points included in the 16S surveys.

**Figure S3.**
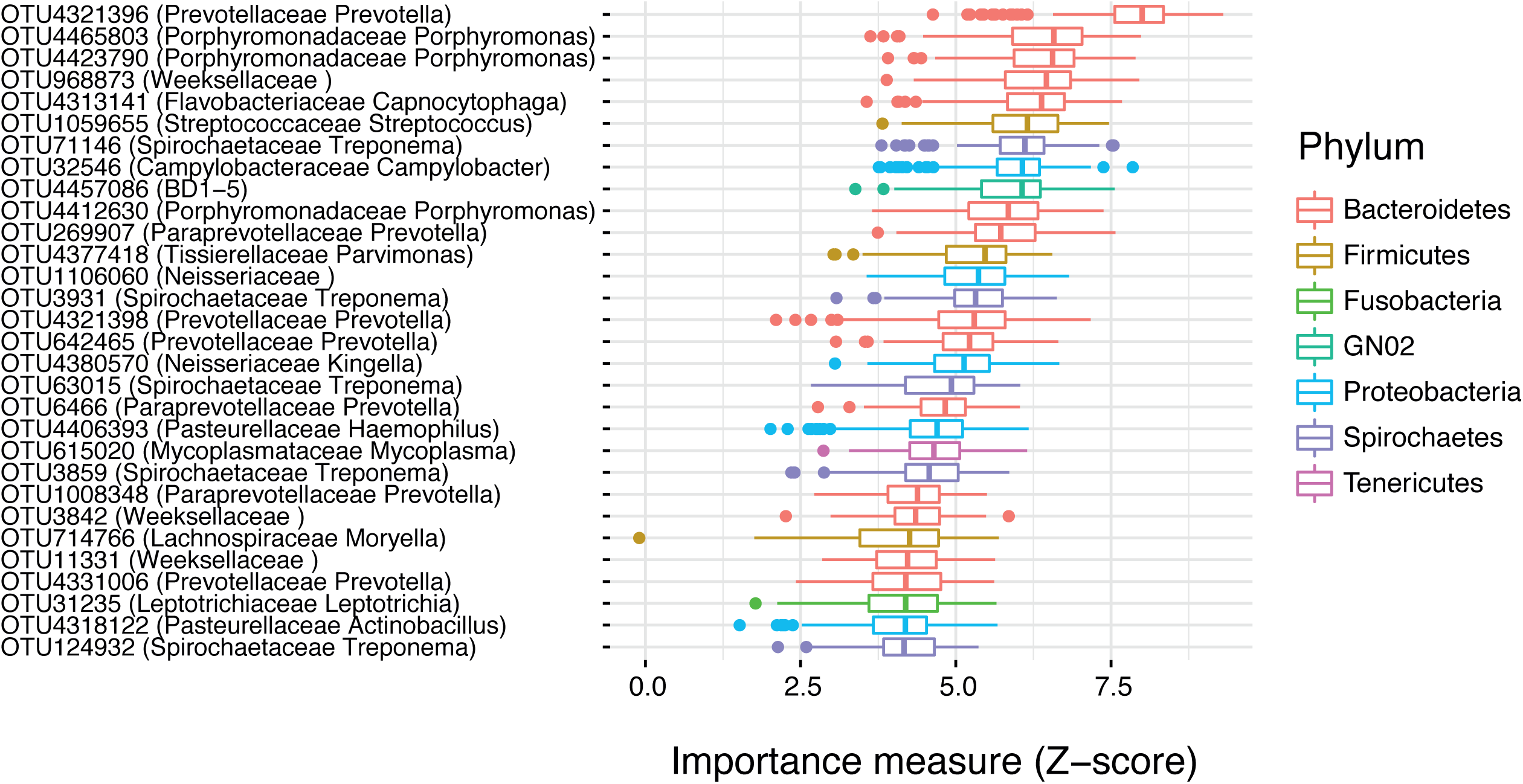
Related to Figure 3. Saliva OTUs differ by SAA group when all 25 individuals assigned to high or low group. We performed random forest analysis using data from all TPs, with all individuals classified into high or low by k-means clustering, such that AMY1M subjects were distributed between the high and low categories. Shown here are the 30 most important OTUs used to predict the SAA group with a random forest classifier, ordered by their Z-score and colored by phyla.

**Figure S4.**
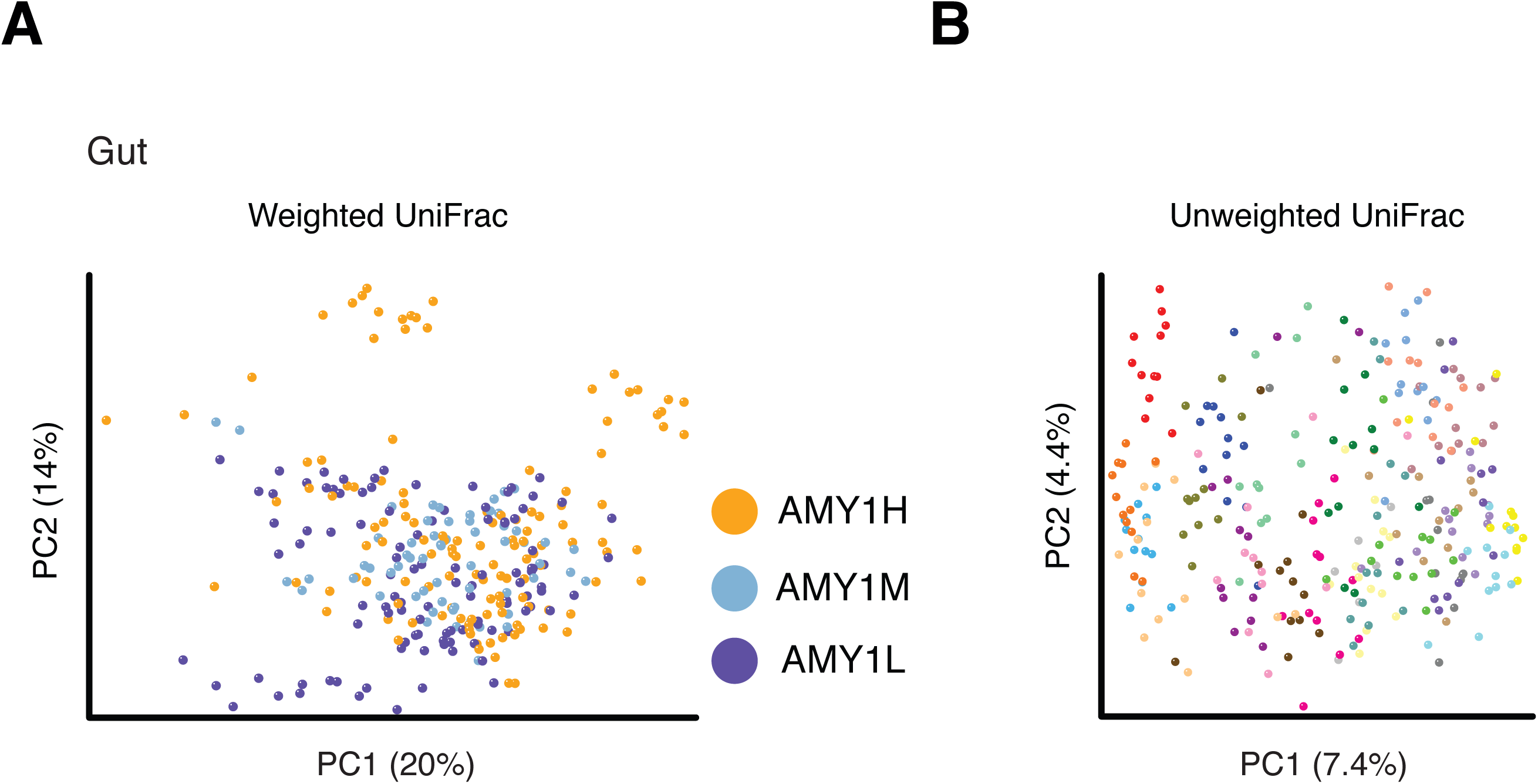
Related to Figure 4. Principal coordinates analysis of stool samples. See legend for Figure S2.

**Figure S5.**
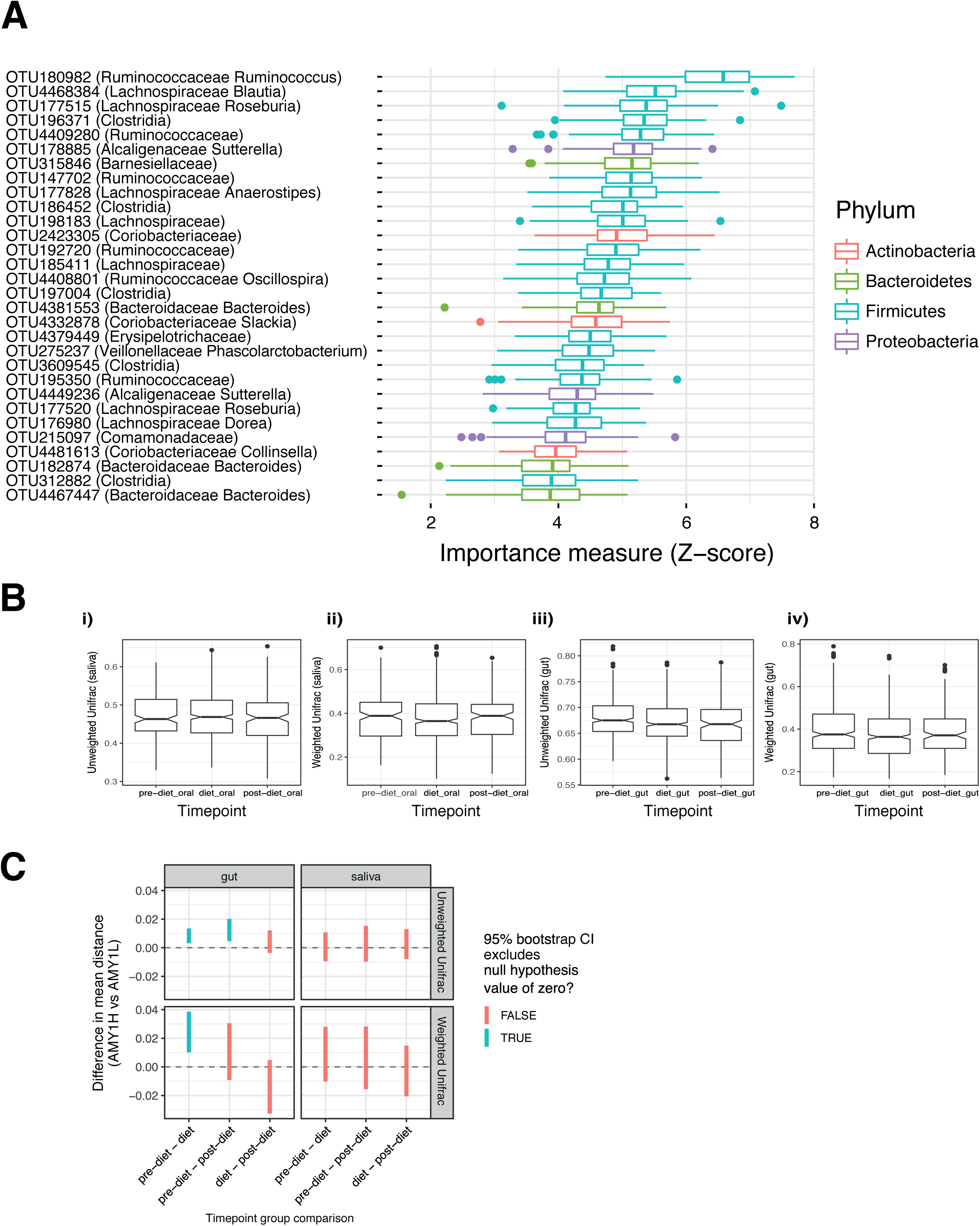
Related to Figure 5. Gut microbiomes differ at OTU level by SAA group and converge on diet. (A) We performed random forest analysis using data from all TPs, with all 25 individuals reclassified into high or low groups by k-means clustering, such that AMY1M subjects were distributed between new high and low categories. Shown here are the top OTUs discriminating between the new high and low groups. (B) The unweighted and weighted UniFrac distances between AMY1H and AMY1L samples between the diet periods. (i) and (ii) are saliva sample data and (iii) and (iv) are gut sample data. (C) For gut and saliva samples, the 95% confidence intervals derived from bootstrap analysis of differences between unweighted and weighted UniFrac distances between AMY1H and AMY1L individuals.

**Figure S6.**
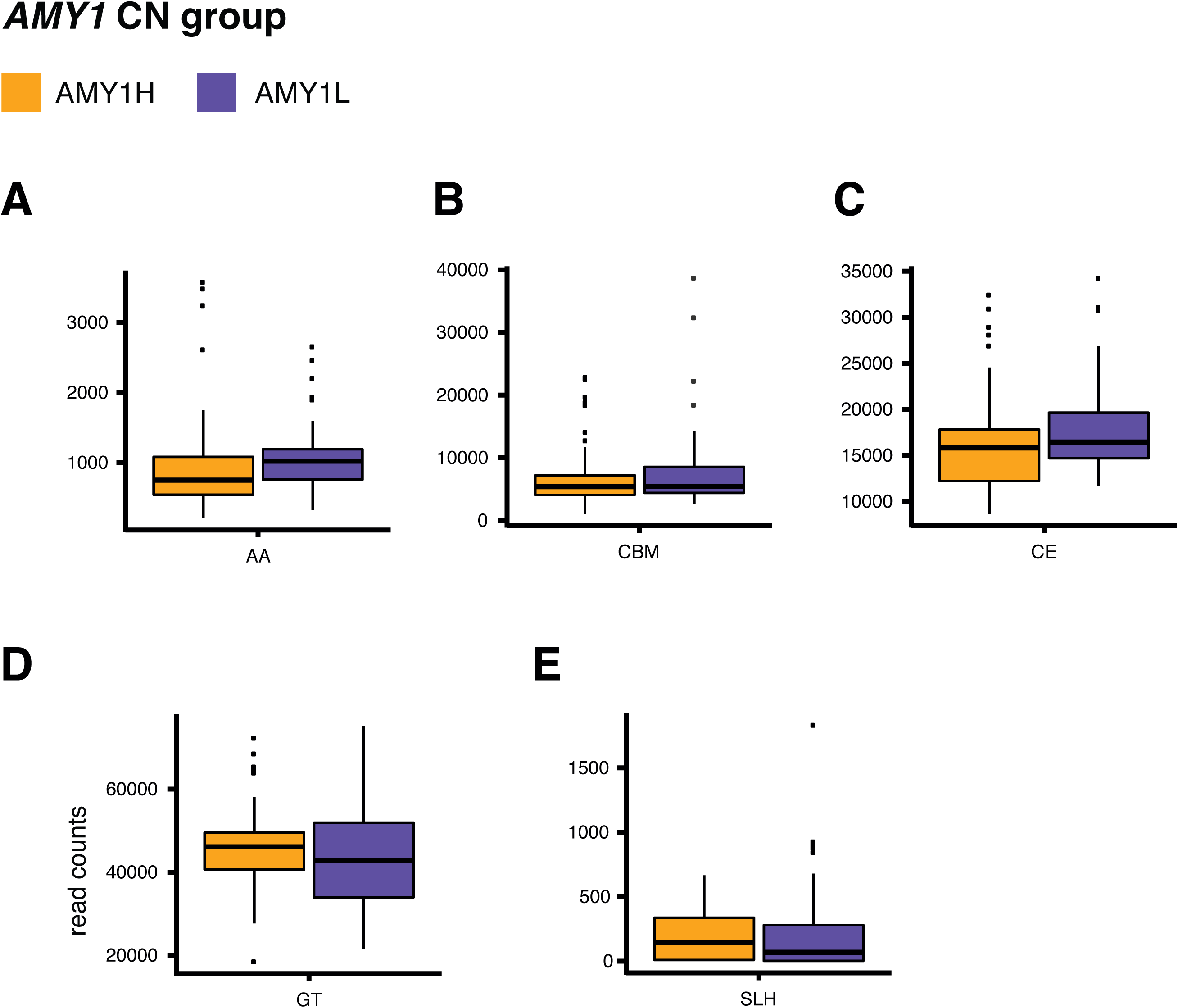
Carbohydrate-Active enZYme classes. Related to Figure 7. Boxplots of the read counts in the AMY1H and AMY1L groups for Carbohydrate-Active enZYme classes, which were not significantly different: (AA) auxiliary activities, (CBM) carbohydrate-binding modules, (CE) carbohydrate esterases, (GT) glycosyltransferases, and (SLH) S-layer homology modules of the cellulosome.

## STAR Methods

### CONTACT FOR REAGENT AND RESOURCE SHARING

Further information and requests for reagents may be directed to, and will be fulfilled by the corresponding author Ruth E. Ley.

### EXPERIMENTAL MODEL AND SUBJECT DETAILS

#### Human subjects

We recruited volunteers affiliated with the Cornell University campus in Ithaca, New York by way of flyers and listservs. We used the following exclusion criteria: BMI ≥ 35 or ≤ 18; age < 18 or > 40; usage within the last six months of amylase inhibitor, systemic antibiotic, corticosteroid, immunosuppressant agent, or probiotic supplement; gastrointestinal disorders. All human-related procedures, sample, and data collection were approved by the Cornell University Institutional Review Board, protocol number 1106002281.

#### Initial *AMY1*-CN screen

We screened 105 individuals on the Cornell University campus for *AMY1*-CN. We collected buccal cells by instructing subjects to swab the inside of their cheek with Epicentre Catch-All Sample Collection Swabs. Genomic DNA was extracted using the Qiagen Gentra Puregene Buccal Cell Kit. qPCR was performed using the same primer sequences for *AMY1* and *TP53* designed by Perry et al. with the following conditions: 2ng of DNA was used in 25μl reactions with Applied Biosystems Power SYBR Green PCR Master Mix on a BioRad MyiQ iCycler Single Color Real-Time PCR Detection System (Perry et al. 2007). The PCR protocol was as follows: initial denaturation at 95ºC for 10 minutes and 40 cycles of 95ºC for 15 seconds followed by 58ºC for 30 seconds. All reactions, including standards, were performed in triplicate. These primers are known to anneal to both human and chimp *AMY1* sequence. Two control sample DNAs with known copy number were run on every plate. The control samples, NS06006: chimp DNA With *AMY1*-CN of 2 and NA18972: human DNA with *AMY1*-CN of 18 (Carpenter et al. 2015; Perry et al. 2007), were purchased from the Coriell Institute for Medical Research NIGMS Human Genetic Cell Repository and NHGRI Sample Repository for Human Genetic Research. To calibrate each subject’s CN, the ratio of *AMY1* to *TP53* levels was used in a line equation created using the ratios from the two control DNA samples resulting in an adjusted copy number value.

#### *AMY1*-CN confirmation

A second cheek swab was taken from each subject to confirm the *AMY1*-CN. Genomic DNA was isolated as described above. We performed qPCR after designing a new set of primers to amplify the *AMY1* paralogs, AMY1-forward: 5’-TGAGAACATTAGGCCACAGCA-3’ and AMY1-reverse: 5’-TGGAAATCATCTCAATGACCTCT-3’. We also designed primers to use *EIF2B2* as a reference gene, EIF2B2-forward: 5’-GCTCAAAGTGCTTGAGGACC-3’, EIF2B2-reverse: 5’-CAAAGCCAAACCCAGACAAT-3’. Primers were at 0.5 μM, and 5ng of DNA template per reaction were used in 10μl reactions with Roche LightCycler 480 SYBR Green I Master mix. Both *AMY1* and *EIF2B2* were run on the same plate, and the PCR program was as follows: initial denaturation at 95ºC for 5 minutes and 40 cycles of 95ºC for 10 seconds followed by 60ºC for 30 seconds on a Roche LightCycler 480 Real-Time PCR Instrument. A standard curve was made using DNA NA10472. Reactions were performed in triplicate, and the control DNAs NS06006 and NA18972 were run on every plate. To calibrate each subject’s CN, the ratio of *AMY1* to *EIF2B2* levels was used in a line equation created using the ratios from the two control DNA samples resulting in an adjusted copy number value.

Digital PCR was performed using Life Technologies Taqman Copy Number Assay Id Hs07226361_cn for the *AMY1* locus and TaqMan Copy Number Reference Assay, RNase P, Human, 4403326 to normalize for total DNA. Reactions were run on a Life Technologies QuantStudio3D Digital PCR system in duplicate.

In statistical analyses (below) we used the mean of the values generated by qPCR and by digital PCR for each subject as the *AMY1*-CN value for that subject.

#### *AMY2* CN determination

We determined the CN of the pancreatic amylase locus, *AMY2*, by performing qPCR with an independent primer pair for each paralog (*AMY2A*, and *AMY2B*), AMY2A-forward: 5’-TGGCGATGGGTTGATATTGCT-3’, AMY2A-reverse: 5’-ACAAGCACAGTGAATTCCGC-3’, AMY2B-forward: 5’-ACTAATGACCTGTGTTATACTTCCT-3’, and AMY2B-reverse: 5’-AGCTGTTACGCACAGTTCCA-3’. We also used the aforementioned primers for *EIF2B2* as a reference gene on the same qPCR plate with each *AMY2* paralog. Primers were at 0.5 μM, and 2ng of DNA template per reaction were used in 10μl reactions with Roche LightCycler 480 SYBR Green I Master mix. *EIF2B2* was run on the same plate with each *AMY2* paralog, and the PCR program was as follows: initial denaturation at 95ºC for 5 minutes and 40 cycles of 95ºC for 10 seconds followed by annealing temperature for 10 seconds and 72ºC for 15 seconds on a Roche LightCycler 480 Real-Time PCR Instrument. Annealing temperature was 58ºC for *AMY2A* and 60ºC for *AMY2B*. Reactions were performed in triplicate.

#### Study design and sample collection

Twenty-five of the aforementioned screened individuals participated in a 4-week study. On day 12 of the study, DEXA scanning was performed on 23 of the subjects using a Hologic DEXA, Model: DISCOVERY-A at Cornell University’s Human Metabolic Research Unit. One subject left the study after two weeks, thus only providing samples for TPs 1 through 6. For the first and fourth weeks, participants were instructed to consume their usual diet and record all food and drink intake with approximate amounts.

During weeks 2 and 3, participants were provided all meals and snacks from a menu designed by a registered dietetic technician. This diet featured healthy meals and snacks and a high-starch food item in every meal. Each day, participants consumed one meal at Cornell University’s Human Metabolic Research Unit dining room in the presence of lab personnel and took two meals away. A researcher from this project took weekday lunches with the participants, and made observations as to the way participants approached the food (e.g., ate very much, very little, avoidance, etc.) Based on the food records and direct observation of behavior at lunch, we are confident that all subjects ate the items provided, albeit in different amounts. On Fridays, participants consumed one meal and took away all meals and snacks packaged for the weekend. Participants were asked to record dietary intake in food diaries during weeks 1 and 4. Dietary checklists were supplied during weeks 2 and 3 when food was provided, and subjects were instructed to record any deviations from the provided menu.

On three days of each of the four weeks, all subjects provided stool and saliva samples for a total of 12 TPs. With few exceptions, all saliva samples were collected on the same day for every subject at each of the twelve TPs. Subjects were instructed to allow saliva to pool in the mouth for three minutes and then express through 5cm drinking straws into a 1.5ml eppendorf tube. Saliva was vortexed and aliquoted into two tubes and chilled on ice until stored at −80ºC within 4 hours of collection. Stool samples were collected during a three-day window for each TP.

#### Dietary intake analysis

Participants recorded dietary intake daily, and afterwards we entered the data into a diet analysis software called SuperTracker. SuperTracker food nutrition data is based on the Food and Nutrient Database for Dietary Studies (FNDDS), and the Food Patterns Equivalents Database (FPED), both from the USDA/ARS Food Surveys Research Group. The software reports categories entitled carbohydrate, dietary fiber, total sugars, and added sugars. Starch is analyzed using the AOAC method 966.11 or 979.10 (2012) or by a polarometric method (The Feedings Stuffs Regulations 1982), but there is not a separate category reported in the SuperTracker output. Total dietary fiber content is determined by enzymatic-gravimetric methods 985.29 or 991.43 of the AOAC (2012). However, the dietary fiber information provided does not distinguish between specific types of fiber including insoluble, soluble, resistant starches and NSPs.

#### Salivary amylase activity

Salivary amylase activity was measured for each saliva sample in triplicate using the SALIMETRICS α-Amylase kinetic enzyme assay kit (cat # 1-1902) as per the instructions with one modification: Instead of 320µl, 300µl of pre-heated substrate was added to the sample. Reactions were performed in triplicate.

#### Stool sample processing

Subjects provided two stool sample aliquots from a single bowel movement in separate tubes and stored them in insulated bags containing frozen ice packs then stored at −80ºC. One of these aliquots was later freeze dried prior to DNA extraction, while the other was saved for use in the amylase activity assay and germfree mouse inoculation studies. At each TP from each subject, one of the stool sample aliquots was freeze dried in a 50ml conical tube and then homogenized by roll-milling in the following manner. After pressing three stainless steel rods (2 different sizes: 9 cm long × 0.9 cm diameter and 9 cm long×0.3 cm diameter) into the freeze dried sample, the 50ml conical tubes were rolled on a Triple Gallon Tumbler (Covington, cat # 253TUM) for 24-48 hours.

#### Stool amylase activity

We used the SALIMETRICS α-Amylase kinetic enzyme assay kit (cat # 1-1902) to measure amylase activity in frozen samples collected during the human studies. We added approximately 150 mg of frozen stool to MOBIO garnet bead tubes with 0.70mm garnet beads (cat # 13123-50). We added enough Salimetrics kit diluent to obtain a concentration of 0.3g stool/ml diluent. Samples were placed in a BioSpec 1001 Mini-Beadbeater-96 for 2 minutes and then centrifuge at 1500 rcf for 15 minutes. We transferred the supernatant into an eppendorf tube and starting with 25μl of undiluted supernatant, performed three serial dilutions up to 1 in 200 using Salimetrics diluent. Then we proceeded with the assay as described above. We noted that 11 subjects (7 AMY1H and 4 AMY1L) had a median FAA< 10 U/g across all TPs.

#### *AMY2* ELISA

We used an ELISA that employs two monoclonal antibodies to human pancreatic amylase as per the instructions (ALPCO, cat # K 6410). As a control, we used purified α-Amylase from *Bacillus licheniformis* (Krackeler Scientific, cat # A3403-500KU) in both the *AMY2* ELISA and amylase activity assays. Amylase activity from *Bacillus licheniformis* was detected in the amylase activity assay, but the ELISA assay specific for *AMY2* did not detect this microbial amylase.

#### Short chain fatty acid analyses on stool samples

Short chain fatty acid quantification was performed by the Metabolomics Core at the University of Michigan using cold extraction of short chain fatty acids, measured by EI GC-MS without derivatization on ~40-60 mg stool from all of the TPs. Short chain fatty acid measurements were normalized to the wet weight of the samples. The short chain fatty acids detected and quantified were acetate, butyrate, propionate, isovalerate, valerate, heptanoate, and hexanoate. This large number of samples had to be run in two batches on different days. There was slight instrumental drift while running the second batch so the LOESS correction method was applied to those data.

#### Microbial DNA extraction, 16S rRNA gene PCR, and sequencing, and QIIME analysis

The saliva samples used in enzyme activity measures, from all TPs except 2, 7, and 10, and all fecal samples, were profiled for microbial community diversity and composition (Table S1). Microbial community DNA was extracted from the freeze dried stool and saliva samples using the MO BIO PowerSoil-htp Soil DNA Isolation Kit (MO BIO Laboratories, Inc., cat # 12955-4), but instead of vortexing, samples were placed in a BioSpec 1001 Mini-Beadbeater-96 for 2 minutes. We used 10-50 ng of sample DNA in duplicate 50 µl PCR reactions with 5 PRIME HotMasterMix and 0.1 µM forward and reverse primers. We amplified the V4 region of 16S using the universal primers 515F and barcoded 806R and the PCR program previously described (Caporaso et al. 2011) but with 25 cycles. We purified amplicons using the Mag-Bind^®^ E-Z Pure Kit (Omega Bio-tek, cat # M1380) and quantified with Invitrogen Quant-iT™ PicoGreen^®^ dsDNA Reagent, and 100 ng of amplicons from each sample were pooled and paired end sequenced (2×250bp) on an Illumina MiSeq instrument. Saliva samples from 3 of 12 TPs were not successfully processed although we did obtain measurements of salivary enzyme activity.

Sequence data were analyzed using the QIIME software package 1.9.0 (Caporaso et al. 2010). Briefly, paired ends were joined using fastq-join, and sequences were demultiplexed and filtered using a Phred quality score threshold of greater than or equal to 25. Open reference OTU picking was performed on all sequence data from oral and gut samples using the uclust method and the August, 2013 Greengenes 16S rRNA Gene Database as reference sequences. We used the QIIME 1.9.0 open reference OTU picking pipeline with all default parameters except the following: max_accepts = 20, max_rejects = 500, stepwords = 20, and word_length = 12. Samples with a sequence count below 10,000 were excluded from downstream analyses. After exclusion, the oral dataset consisted of 216 samples (sequencing was performed on 9 of the 12 TPs collected) yielding 16,030,493 sequences with a median sequence count of 72,107. The fecal data set included 293 samples with a total of 16,421,608 sequences and a median sequence count of 55,165 sequences per sample.

We calculated beta diversity using the unweighted and weighted UniFrac metrics on an OTU table containing 11,146 and 20,133 sequences per sample for the oral and gut datasets, respectively, and principal coordinates analysis on the distance matrices (Lozupone et al. 2007). We assessed alpha diversity using Chao 1, Observed Species, Faith’s phylogenetic diversity, and Shannon’s Index (Magurran 2004; Southwood and Henderson 2009) by calculating means from 100 iterations using a rarefaction of 11,146 and 16,848 sequences per sample for the oral and gut datasets, respectively.

#### Metagenomics sample preparation, sequencing, and analysis

We performed metagenomic analysis on sequences generated from genomic microbial DNA obtained during the DNA extraction method detailed above using the freeze dried stool samples collected at TPs 3, 4, 6, 7, 9, and 10 (Table S1). We prepared metagenomic libraries using 1 ng of DNA input per sample into a Nextera XT DNA Sample Preparation Index Kit as per the instructions (Illumina, Inc., cat # FC-131-1096). After purification with Agencourt AMPure XP beads (Beckman Coulter, cat # A63882), samples were normalized and pooled with 20 samples per pool. Size selection was performed on the pools using BluePippin (Sage Sciences, cat # BDF1510) to restrict fragment sizes between 300 to 650 bp. Pools were run on an Illumina HiSeq3000 with 2×300 bp paired end sequencing for a sequencing depth of 14 ± 3.0 Gb (median ± standard deviation).

#### Germfree mouse transfer experiments

All mouse-related protocols, sample, and data collection were approved by the Cornell University Institutional Animal Care and Use Committee, protocol number 2010-0065. We inoculated germfree Swiss Webster adult male mice between 4-6.5 weeks of age with stool samples collected at 5 TPs (3, 4, 7, 9, and 10) during the human study (Table S1). Each mouse experiment used stool samples from all 9 AMY1L donors. TP 3 had 11 AMY1H donors while all other TPs used 10.

At the beginning of each experiment, mice were weight-ranked and then assigned alternately to an AMY1L or AMY1H donor to ensure that the mean weight on the day of inoculation was not different between the 2 groups. Each mouse was orally gavaged with 200 µl stool suspension from one human subject and single housed. Stool suspension was prepared in a Coy anaerobic chamber. Approximately 500 mg frozen stool was solubilized in 10ml of anaerobic PBS that contained 2 mM DTT as a reducing agent and vortexed at 5 minute intervals until no soluble clumps were visible. After inoculation, mice were maintained on autoclaved water and autoclaved Teklad diet 7017, NIH-31 (Harlan Laboratories) and kept under a 12-hour light/dark cycle for 33-40 days (TP 3: 40 days; TP 4: 35 days; TP 7: 35 days; TP 9: 33 days; and TP 10: 35 days.) Mouse weight and chow consumption were recorded weekly. At the end of each experiment, mice were sacrificed and DEXA scanned (Lunar PIXImus Mouse, GE Medical Systems, Waukesha,WI) to measure adiposity.

## QUANTIFICATION AND STATISTICAL ANALYSIS

We excluded the medium group from all statistical analyses that include *AMY1* group as a covariate but included them in all analyses with *AMY1*-CN as a covariate. We used logarithm, square root or rank transformation as needed to better meet model requirements of homogeneous variance and normality of residuals, ε. All linear mixed models are described with the notation used in the statistical package lme4 in R, version 3.1.2 (Bates et al. 2014). The Scikit-Learn library in Python was used in the machine learning based modeling.

### Macronutrient intake analysis

Dietary records from all subjects were manually entered into the nutritional analysis software SuperTracker, which produces nutrient reports that include estimated percentages of macro- and micronutrient content in the food items entered by the user. We fit linear mixed models using each macronutrient percentage as the response variable to determine whether or not dietary intake differed between the *AMY1* CN groups over time. We analyzed two separate models that either included study day or whether or not diet was being provided on that day:

1. *y* ~ **AMY1*C*NG** + **DAY** + **AMY1CNG**: **DAY** + (1|**SUBJECT**) + ε
2. *y* ~ **AMY1CNG** + **DIET** + **AMY1CNG**: **DIET** + (1|**SUBJECT**) + ε

***y*** is the macronutrient percentage. Fixed effects included *AMY1* CN group (**AMY1CNG**) and study day (**DAY**) or whether or not diet was being provided on that day (**DIET**). We also included a random effects term for repeat sampling of subjects (1|**SUBJECT**).

### Salivary amylase activity, linear mixed models using either group or copy number

We analyzed two separate models that included either *AMY1* CN or *AMY1* CN group as a predictor:

1. *y* ~ **AMY1CNG** + **TP** + **AMY1CNG**: **TP** + (1|**SUBJECT**) + ε
2. *y* ~ **AMY1CN** + **TP** + **AMY1CN**: **TP** + (1|**SUBJECT**) + ε

***y*** is the salivary amylase activity. Fixed effects included *AMY1* CN group (**AMY1CNG**) or *AMY1* CN (**AMY1CN**; included subjects in the medium CN group), and TP (**TP**). We also included a random effects term for repeat sampling of subjects (1|**SUBJECT**).

Neither the interaction between *AMY1* CN group and TP nor *AMY1* CN and TP is significant and nor is the effect of TP significant in either model. The effect of both *AMY1* CN (p = 2.1×0^−5^) and *AMY1* CN group (p = 1.9×0^−4^) are significant based on an F-test with a Satterthwaite approximation.

### Effects of diet on distances between AMY1H and AMY1L individuals

For 16S data, we compared Unifrac distances (both weighted and weighted) between individuals in high and low *AMY1* groups during 3 time intervals: pre-diet, on the diet, and post-diet. For shotgun metagenomic sequencing, we compared Bray Curtis distances between individuals in high and low *AMY1* groups using the gene family raw counts and only had data for TPs during the time intervals: pre-diet and on the diet. We calculated the non-parametric bootstrap confidence intervals for the difference in population means between the Bray-Curtis distances (just AMY1L versus AMY1H distance values) for pre-diet and during-diet time points. We determined the 95% bootstrap CIs based on 1000 permutations. This approach accounts for non-independence issues caused by repeat sampling from individuals.

### Alpha diversity in stool and saliva, linear mixed models using either group or CN

We used a linear mixed model to assess the effect of *AMY1* CN or *AMY1* CN group on alpha diversity:

1. *y* ~ **AMY1CN** + **TP** + **AMY1CN**: **TP** + (1|**SUBJECT**) + ε
2. *y* ~ **AMY1CNG** + **TP** + **AMY1CNG**: **TP** + (1|**SUBJECT**) + ε

***y*** is the alpha diversity metric. Fixed effects included *AMY1* CN group (**AMY1CNG**; excluded medium group) or *AMY1* CN (**AMY1CN**; included subjects in the medium CN group) and TP (**TP**). We also included a random effects term for repeat sampling of subjects (1|**SUBJECT**).

Whenever the interaction term of the linear mixed model was significant, we identified the affected TPs by performing post-hoc pairwise comparisons between the TPs using Tukey’s HSD method to adjust for multiple comparisons.

### OTU relative abundances between *AMY1* CN groups

We used a bivariate model called Harvest (Bar, Booth, and Wells 2014) to identify OTUs with differential means or variances in relative abundance between the AMY1H and AMY1L groups at each TP separately. For this analysis we omitted OTUs not present in at least half of the samples in either the AMY1L or the AMY1H group at the TP being considered. We adjusted p values using the Benjamini-Hochberg procedure to account for all OTUs tested at a given TP.

### Estimation of *AMY1* CN distribution in British population

Included in the British genotype data were 7 of the 10 SNPs, rs 6696797, rs 10881197, rs 1999478, rs 11185098, rs 1930212, rs 1566154, and rs 1330403, previously correlated with *AMY1* CN (Usher et al. 2015). Only one randomly chosen twin per pair was included in our analysis. After excluding anyone with a BMI outside of that used to screen our Cornell population, we had data for 994 British subjects. Using the change in copy number values determined for the GoT2D cohort of 2,863 Europeans, we calculated the sum of the change in copy number values corresponding to each person’s 7 alleles. We then selected only the tail ends of the distribution to include the lowest 5% and highest 5% of individuals for group sizes of 50 each.

### Stool amylase activity, linear mixed models using either group or CN

We used a linear mixed model to assess the effect of *AMY1* CN, *AMY2* CN, or *AMY1* CN group on stool amylase activity using the following models:

1. *y* ~ **AMY2CN** + **TP** + **AMY2CN**: **TP** + (1|**SUBJECT**) + ε
2. *y* ~ **AMY1CN** + **TP** + **AMY1CN**: **TP** + (1|**SUBJECT**) + ε
3. *y* ~ **AMY1CNG** + **TP** + **AMY1CNG**: **TP** + (1|**SUBJECT**) + ε

***y*** is the stool amylase activity. Fixed effects included *AMY1* CN (**AMY1CN**; included subjects in the medium CN group), *AMY2* CN (**AMY2CN**; included subjects in the medium CN group), or *AMY1* CN group (**AMY1CNG**; excluded medium group), and TP (**TP**). We also included a random effects term for repeat sampling of subjects (1|**SUBJECT**).

In model [3] we determined that stool amylase activity at TP 6 is significantly greater than TP 12 by performing post-hoc pairwise comparisons between the TPs using Tukey’s HSD method to adjust for multiple comparisons.

### Metagenomics analysis

For metagenomic read quality control, we used skewer v0.2.2 (Jiang et al. 2014) to trim the 3’ ends until quality scores reached ≥15, and end-trimmed reads <100 bp were removed. skewer v0.2.2was also used to remove Illumina adapter contamination, and bbmap v37.78 was used to filter out human host genome reads. Post-QC reads were subsampled at 20,000,000 paired-end reads per sample to normalize for sequencing depth and reduce downstream processing time. We used the HMP Unified Metabolic Analysis Network (HUMAnN2 v0.11.1) pipeline to classify the reads against the ChocoPhlAn and UniRef90 databases (Abubucker et al. 2012). For our statistical analyses, we used the gene families file with the abundances normalized to reads per kilobase (RPKs). However, DESeq2, the software that we used to identify gene families with differential abundances between AMY1 groups, requires integer values, or counts that have not been normalized with respect to library size, as a requirement of the statistical model because the DESeq2 software adjusts for differences in library size internally (Love et al. 2014). To accommodate this requirement, we multiplied the RPKs reported by HUManN2 by the gene lengths and total number of reads in order to obtain integers unadjusted for library size. The gene families file produced by HUMAnN2 is stratified; for each gene family there are one or more rows with the first row being the total number of reads assigned to that gene family and the additional rows corresponding to the number of reads assigned to each of the different taxonomy, i.e. species, when known. Therefore, the raw data contains entries that are not independent. We removed the entry reporting the sum of the mappings assigned to the gene family prior to analysis and kept the mappings to species including unclassified. We also filtered out gene families not present in at least half of the samples in either the AMY1L or the AMY1H group in the dataset. Then we identified the differentially abundant gene families at each TP using DESeq2 and used the log2 fold change between AMY1H and AMY1L for each gene family to create a heatmap. We adjusted p values using the Benjamini-Hochberg procedure to account for all gene families tested at a given TP and displayed gene families with BH-adjusted p values < 0.01. Furthermore, the heatmap includes only gene families with assigned taxonomy and is sorted by taxonomy. When a gene family is not significant at a TP, the corresponding heatmap cell is colored gray. The heatmap in Figure 6 was created using the software iTOL with gene families ordered by taxonomy.

### Carbohydrate-Active enZYmes analysis

We used hmmscan to query the gene families with HMMs from dbCAN (release 6.0), and used an e-value cutoff of 1e-18 to positively identify CAZYmes.

We used linear mixed models to determine whether the number of read counts from any of the CAZYme classes differed between the AMY1H and AMY1L groups.

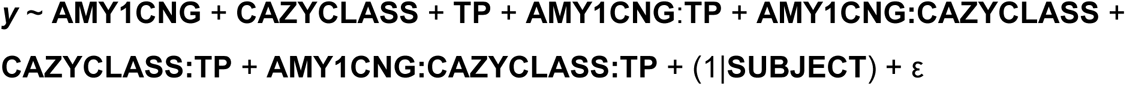

**y** represents the number of read counts, and fixed effects are *AMY1* CN group (**AMY1CNG**), TP (**TP**), and CAZYme class (**CAZYCLASS**). We also included a random effects term for repeat sampling of subjects (1|**SUBJECT**).

### Determination of SAA groups

Each of the 25 participants were labeled as a member of either SAA-H or SAA-L group. These labels were assigned using the KMeans module, which implements the k-means algorithm, and was parameterized to identify two clusters within the mean salivary amylase activity measurements.

### Identification of OTUs distinguishing SAA groups in saliva and stool

Using phyloseq v1.22.3, we filtered out OTUs with an average relative abundance below 0.001%. This screening produced two data sets composed of 216 salivary and 283 fecal samples with 672 and 900 taxa, respectively. Samples were labelled according to the salivary amylase activity level assigned to its subject of origin. For the machine learning analysis, the OTUs were considered features, and the salivary amylase activity level (high or low) as the response variable. We used caret v6.0 to create a partition with 80% of the samples for training, and the remaining 20% for testing purposes. Then, we executed a random forest model on the training data set using the randomForest v4.6 package, adjusting the number of trees to 200 and enabling the flag to calculate the feature importances. Feature selection was done using Boruta v5.2.0 with default parameters on the training data set (after excluding or not OTU abundance measures coming from subjects originally placed in the *Medium* group). All the OTUs confirmed as important or tentative were treated as relevant, resulting in 113 for saliva and 301 for feces. To examine the predictive power of the model, we used the caret package and trained two random forest models with default parameters, one removing the relevant OTUs, and the other with only same relevant OTUs.

### Prediction of salivary amylase activity with short chain fatty acid measurements in stool

The dataset was divided in two subsets, one for the training process with 266 records, and another with 54 records for testing purposes. Then, a RandomForestClassifier object was configured to use 250 estimators, and automatically adjusted their weights with respect to the frequency of SAA group. Once the classifier was trained, its predictive performance was assessed using the following statistics: the F1 score, which represents the harmonic average of the precision and recall, and the Matthews correlation coefficient. To visualize the classification results, we generated a confusion matrix and a receiver operating characteristic (ROC) curve. Finally, the importance of features was calculated from the Gini impurity criteria used by the classifier to evaluate the quality of the resulting decision trees.

For this linear mixed model, the salivary amylase activity was transformed to better fit a normal distribution using the transformTukey() function from the rcompanion package. Based on the contributions of SCFA to the prediction of the salivary groups: (1) heptanoate, isovalerate and hexanoate were excluded from the analysis and (2) geometric means of grouped SCFA were calculated on the concentrations and on the percentages. The groups were the following: {total concentration and butyrate}, {valerate, propionate, acetate}, {all together}. The subjects and TPs were included as random effects (due to the fact that they were not taken into account in the random forest analysis). The best model was chosen by a forward method until all predictors were significant and the p-values from the final model were corrected using the Benjamini-Hochberg method. The final model is the following:

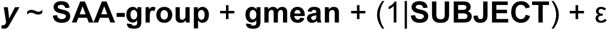

with **y** the salivary amylase activity and **gmean** the geometric mean of the total concentration of SCFA and of the concentrations of butyrate, valerate, propionate and acetate:

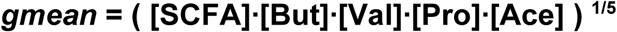

### Assessment of adiposity in mice

We used a linear mixed model to determine whether adiposity differed between AMY1H and AMY1L microbiome recipients using the following equation:

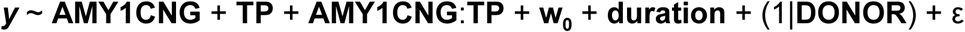

**y** is the percent fat determined by DEXA, and effect terms include *AMY1* CN group, time point, weight on the day of inoculation, and duration or length of experiment (**AMY1CNG**, **TP**, **w**_**0**_, **duration**), and we included a random term for human donor. We identified the affected TPs by performing post-hoc pairwise comparisons between the TPs using Tukey’s HSD method to adjust for multiple comparisons.

